# A human brain vascular atlas reveals diverse cell mediators of Alzheimer’s disease risk

**DOI:** 10.1101/2021.04.26.441262

**Authors:** Andrew C. Yang, Ryan T. Vest, Fabian Kern, Davis P. Lee, Christina A. Maat, Patricia M. Losada, Michelle B. Chen, Maayan Agam, Nicholas Schaum, Nathalie Khoury, Kruti Calcuttawala, Róbert Pálovics, Andrew Shin, Elizabeth Y. Wang, Jian Luo, David Gate, Julie A. Siegenthaler, M. Windy McNerney, Andreas Keller, Tony Wyss-Coray

## Abstract

The human brain vasculature is of vast medical importance: its dysfunction causes disability and death, and the specialized structure it forms—the blood-brain barrier—impedes treatment of nearly all brain disorders. Yet, no molecular atlas of the human brain vasculature exists. Here, we develop Vessel Isolation and Nuclei Extraction for Sequencing (VINE-seq) to profile the major human brain vascular and perivascular cell types through 143,793 single-nucleus transcriptomes from 25 hippocampus and cortex samples of 17 control and Alzheimer’s disease (AD) patients. We identify brain region-enriched pathways and genes divergent between humans and mice, including those involved in disease. We describe the principles of human arteriovenous organization, recapitulating a gradual endothelial and punctuated mural cell continuum; but discover that many zonation and cell-type markers differ between species. We discover two subtypes of human pericytes, marked by solute transport and extracellular matrix (ECM) organization; and define perivascular versus meningeal fibroblast specialization. In AD, we observe a selective vulnerability of ECM-maintaining pericytes and gene expression patterns implicating dysregulated blood flow. With an expanded survey of brain cell types, we find that 30 of the top 45 AD GWAS genes are expressed in the human brain vasculature, confirmed *in situ*. Vascular GWAS genes map to endothelial protein transport, adaptive immune, and ECM pathways. Many are microglia-specific in mice, suggesting an evolutionary transfer of AD risk to human vascular cells. Our work unravels the molecular basis of the human brain vasculature, informing our understanding of overall brain health, disease, and therapy.

## Main Text

Brain health depends on brain vascular health. The brain is one of the most highly perfused organs in the body, necessary to meet its unique metabolic needs^1^. Brain vascular dysfunction is a major contributor to stroke, the second leading cause of death worldwide^2–4^. Dysfunction causes serious long-term disability: vascular-specific genes are mutated in congenital neurological disorders^5, 6^ and age-related vascular impairments are increasingly appreciated in neurodegenerative disease^7–11^. The brain vasculature moreover forms a special structure, the blood-brain barrier (BBB), that mediates selective movement of molecules between the blood and the brain^12–15^. While necessary for optimal neuronal function^16, 17^, the BBB frustrates the pharmacological treatment of nearly all brain disorders^15, 18, 19^, and extensive efforts are underway to identify targets on the BBB for enhanced drug delivery^20–22^. These brain vascular properties arise from a complex ecosystem of cells and their interactions^17, 23, 24^: endothelial cells, adjacent mural smooth muscle cells and pericytes, perivascular immune cells, and surrounding astrocytes that differ across brain regions and vary along an arteriovenous gradient^14, 25, 26^. Heterogeneity along this gradient produces functionally segmented circulatory, metabolic, and permeability properties^6, 27^.

Recent studies have characterized the cellular heterogeneity of the human brain in health and disease using single-nucleus RNA sequencing (snRNA-seq)^28–33^. They have elucidated cell type-specific perturbations in multiple sclerosis, autism, and Alzheimer’s disease; pinpointed which cell types express risk genes identified in genome-wide association studies (GWAS); and nominated biological pathways for further study. Yet, though vascular cell density^34, 35^ is estimated at 70,000 cells/mm^3^ (approaching total glia density^34, 36^), such studies, to our knowledge, have mostly lost these cells during the isolation process for unknown reasons. Pioneering work has profiled the mouse brain vasculature^37–42^, but it remains unclear how conserved these findings are in humans, given the approximately 96 million years of evolutionary divergence^43^. Indeed, recent studies have documented species-specific pathways in microglia, notably in disease GWAS loci^44^; and recent attempts to advance brain-penetrant AAVs into the clinic have stalled because of mouse-specific expression of the cognate endothelial receptor LY6A^45–47^. Moreover, mouse brain vascular sequencing studies so far have been limited to the non-diseased setting and without regard to brain region heterogeneity.

Given the scientific, medical, and pharmacological importance of the human brain vasculature, we set out to systematically characterize the principal vascular cell types in both the hippocampus and cortex of control and AD patients.

### Cells of the human brain vasculature

We hypothesized that unlike parenchymal nuclei, vascular cells and nuclei remain entombed in the basement membranes of blood vessels after typical dounce homogenization and processing of frozen brain tissue for snRNA-seq^28–33, 48–52^. Such vessel fragments are caught on strainers prior to droplet capture, and vessel fragments that do pass through can yield doublet or hybrid nuclei resulting in unreliable or artificial clusters upon analysis. Thus, we set out to develop methods to first physically isolate brain vessels and then extract discrete nuclei from them. Specifically, after density centrifugation^53^ and strainer capture^54^, we tested various enzymatic (e.g., papain, collagenase, trypsin), chemical (e.g., osmolarity, detergents), and physical (e.g., sonication, TissueRuptor) approaches to liberate nuclei. Nearly all resulted in nuclei damage or nuclei devoid of RNA reads. We finally found success adapting a gentle protocol for splenocyte isolation^55^ (Methods)—and combined it with extensive sucrose and FACS-based cleanup to ensure high-quality data (Fig. 1a, Supplemental Fig. 1).

**Figure 1.**
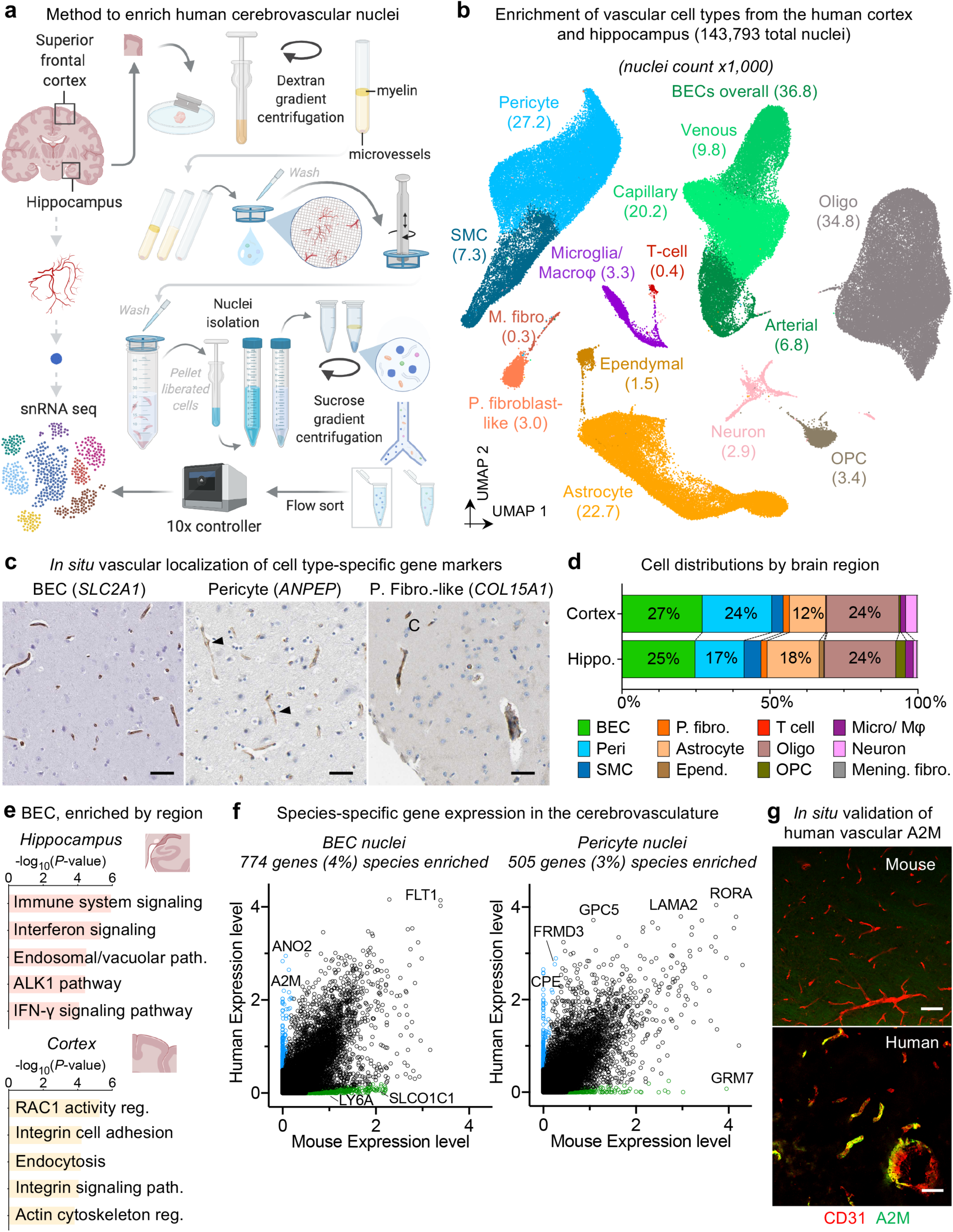
**Cells of the human brain vasculature.** **a**, Method optimized to enriched cerebrovascular nuclei from post-mortem human brain samples. **b**, Uniform Manifold Approximation and Projection (UMAP) of 143,793 nuclei captured from 25 human hippocampus and superior frontal cortex samples across 17 patients, colored by cell type and labeled with the number of nuclei. **c**, Immunohistochemical validation of cell type-specific gene markers from (**b**) for brain endothelial cells (BECs), pericytes, and perivascular fibroblast-like cells. Arrowhead (middle) identifies staining of thin-strand pericyte morphology. C (right) indicates capillary staining. Scale bar = 50 microns. Images from the Human Protein Atlas (http://www.proteinatlas.org)^75, 140^. **d**, Distribution of cell populations, identified from nuclei isolated from the hippocampus and frontal superior cortex, after sub-clustering analysis. Color code corresponds to (**b**). **e**, Enriched biological pathways in BECs from the hippocampus compared to the superior frontal cortex, and vice versa, in control patient samples. **f**, Scatter plot depicting mRNA expression levels (logCPM) of mouse and human genes with one-to-one orthologs in BECs (left) and pericytes (right), highlighting significantly differentially expressed genes in human (blue) and mouse microglia (green) (>10-fold difference, minimum 0.5 log_2_CPM expression). **g**, Immunohistochemical validation of A2M protein localized specifically in the human but not mouse vasculature. Scale bar = 50 microns.

With our new method, which we call VINE-seq (Vessel Isolation and Nuclei Extraction for Sequencing), we processed 25 samples: the hippocampus of 9 AD and 8 age- and sex-matched controls, as well as the superior frontal cortex from a subset of 8 patients (4 samples per group, Supplemental Table 1). Samples included a range of *APOE* genotypes (E3/3, E3/4, E4/4). After quality-control (Methods), we obtained 143,793 single nucleus transcriptomes. Visualization in uniform manifold approximation and projection (UMAP) space separated nuclei into distinct clusters, which we mapped to 15 major cell types (Fig. 1b), including all known vascular and perivascular cell types, many not captured before from human brains: endothelial cells (arterial, capillary, venous), smooth muscle cells, pericytes, astrocytes, perivascular macrophages, T cells, and both perivascular and meningeal fibroblasts. The number of cerebrovascular nuclei captured here exceed those in the literature by at least several hundred-fold (Fig. 1b, Supplemental Fig. 2). Canonical markers used to identify cell types (Supplemental Fig. 3-4, Supplemental Table 2) were validated for their predicted vascular localization *in situ* (Fig. 1c). Expression levels for each gene across cell types are available to browse at https://twc-stanford.shinyapps.io/human_bbb.

The distributions of cell types differed between the hippocampus and frontal cortex (Fig. 1d). Astrocyte and oligodendrocyte progenitor cell (OPC) frequencies were higher in the hippocampus, recapitulating prior cell density studies^34, 56^. Amongst all cell types, astrocyte transcriptional identity was the most influenced by brain region, forming distinct hippocampus- and cortex-enriched cell subclusters (Supplemental Fig. 4d-g). Pericytes, critical for regulating blood-brain barrier (BBB) function^15, 23, 24^, were reduced already in control hippocampi relative to cortices. Each vascular cell type exhibited brain region-specific enrichments in genes and pathways (Supplemental Table 3). For example, hippocampal endothelial cells demonstrated greater baseline inflammation, such as IFN-γ signaling, than those in the cortex (Fig. 1e). Such inflammatory signaling has recently been described to inhibit hippocampal neurogenesis^57–59^, and together with the aforementioned pericyte loss, provides a molecular hypothesis for the particular susceptibility of the hippocampal vasculature to dysfunction in both aging^8, 54^ and AD^55^.

We next compared nuclei transcriptomes between human and mouse endothelial cells and pericytes. Using a strict cutoff (>10x difference, logCPM > 0.5, Supplemental Table 4), we found hundreds of species-enriched genes (Fig. 1f). These include the known mouse-specific endothelial anion transporter *Slco1c1*^60^ and AAV PHP.eB receptor *Ly6a*^45–47^. These also include disease-related genes such as *A2M* and *CASS4*, implicated in β-amyloid processing (Fig. 1g)^61–64^. Several small molecule transporters varied, suggesting species differences in brain metabolism. For instance, the GABA transporter *SLC6A12* is enriched in human over mouse pericytes, with implications for GABAergic neurotransmission and associated diseases like epilepsy^65^. We confirmed the vascular localization of *SLC6A12* and other human-enriched genes in human brain tissue at the protein level (Fig. 1g, Supplemental Fig. 5). Several genes of high pharmacological importance mediating small molecule and protein BBB transport vary between species (Supplemental Fig. 6). Finally, this dataset enables study of diseases that involve the human brain vasculature, such as genes relevant to SARS-CoV-2 neuroinvasion^66, 67^, neurotoxicities associated with cancer immunotherapies^68^ (e.g., no *CD19* expression in human adult brain pericytes), and the cell type etiology of ALS^69^ (Supplemental Fig. 7). Together, the VINE-seq method introduced here opens the human brain vasculature for molecular study and provides an important data resource for interrogating its diverse cell types.

### Organizing principles of human brain endothelial and mural cells

With our capture of large numbers of vascular nuclei (>30x mouse^37^, >200x human^28, 29^ prior studies), we sought to comprehensively characterize the molecular basis of endothelial and mural cell organization along the human brain arteriovenous axis. Cellular and molecular changes along this axis have been referred to as zonation^15, 37, 41, 70, 71^. Beginning with endothelial cells, a UMAP representation resolved the 36,825 captured nuclei into the known vessel segments: arterial and venous clusters were located at opposite ends, separated by a major capillary cluster (Fig. 2a, Supplemental Fig. 8a-b). These clusters were defined by established zonation markers, such as arterial *VEGFC* and *ALPL*; capillary *MFSD2A* and *SLC7A5*; and venous *IL1R1* and *NR2F2*^37, 41^. While capillaries make up the vast majority (∼90%)^27, 72^ of the endothelium, our method facilitated robust capture of rarer arterial (at 19%) and venous (at 27%) endothelial cells, likely either because of more efficient strainer retention or nuclei liberation. We also noticed a small endothelial cluster (571 nuclei, ∼0.1%) outside conventional arteriovenous zonation. This cluster expressed genes characteristic of ‘tip’ cells (e.g., *PLAUR* and *LAMB1*)^73^ as well as ‘proteostatic’ heat shock proteins^37^.

**Figure 2.**
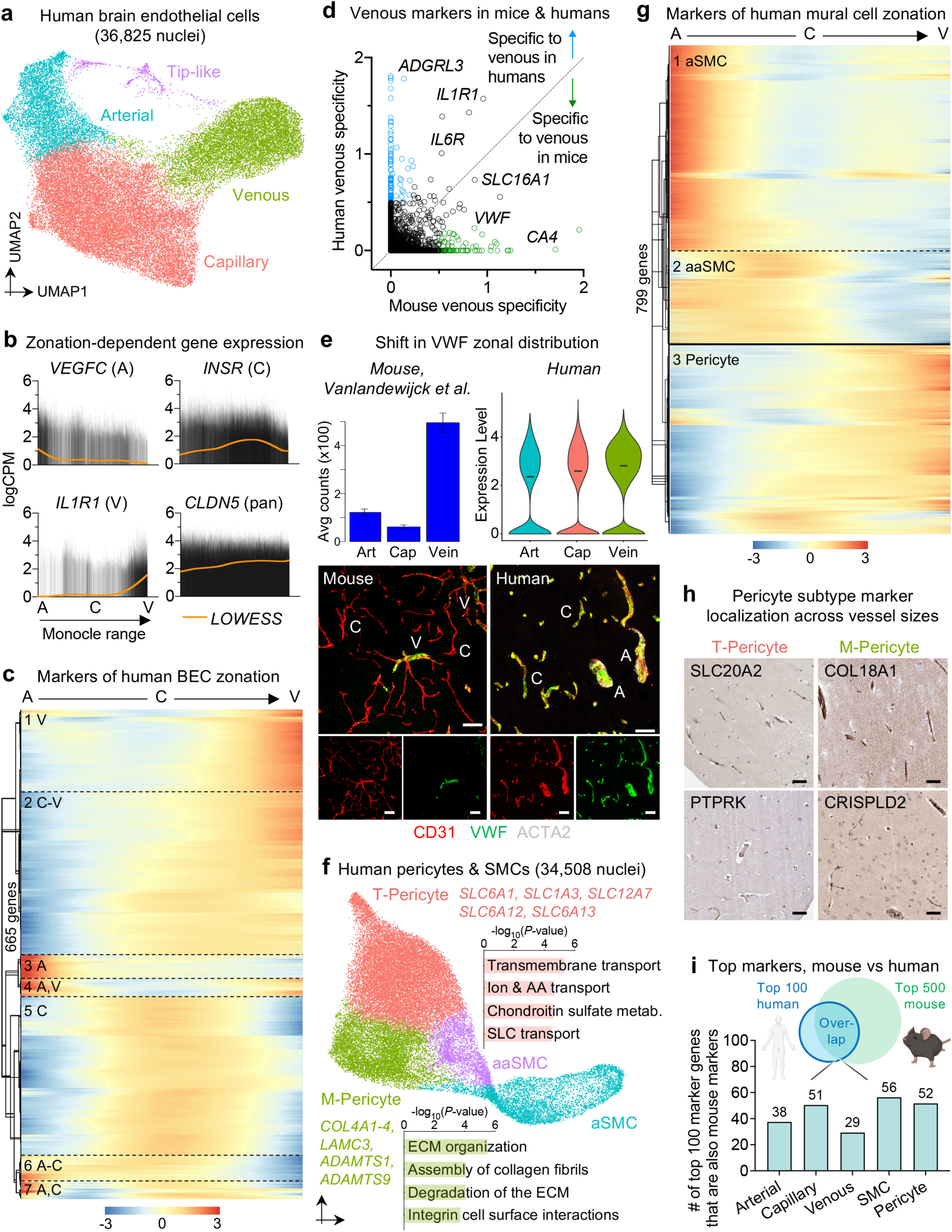
**Organizing principles of human brain endothelial and mural cells.** **a**, UMAP of 36,825 human brain endothelial cell (BEC) nuclei, colored by zonation. **b**, Zonal expression of transcripts across human BECs sorted by Monocle pseudotime. LOWESS local regression line (orange) and density of black lines (counts) correspond with average expression levels. A = arterial, C = capillary, and V = venous. **c**, Heatmap of zonation-dependent gene expression in human BECs. **d**, Scatter plot depicting the specificity of transcripts for venous BECs in mouse^71, 90^ compared to humans. Venous specificity score = avg(logFC(vein/cap), logFC(vein/art)). For example, *VWF* is predicted to be more specific to venous BECs in mice than it is in humans. See Supplemental Figure 9 for arterial and capillary specificity plots. **e**, Immunohistochemical validation of VWF specificity to venous BECs in mice but not in humans. Comparison of *VWF* gene expression in mice and human BECs (top) and corresponding VWF protein staining (bottom). Scale bar = 50 microns. **f**, UMAP of 34,508 human pericyte and smooth muscle cell nuclei, colored by cell subtype. Enriched biological pathways from directly comparing amongst the pericyte and SMC populations. aSMC = arterial smooth muscle cell (aSMC), aaSMC = arteriole SMCs, T-Pericyte = solute transport pericytes, and M-Pericyte = Extracellular matrix regulating pericytes. **g**, As in (**c**) but for pericytes and smooth muscle cells. Solid line delineating aaSMC/aSMCs from pericytes reflects an abrupt transcriptomic transition. **h**, Immunohistochemical localization of T- and M-pericyte markers in both large and small diameter vessels, suggesting pericyte subtypes do not represent a pericyte-vSMC split. Scale bar = 50 microns. Images from the Human Protein Atlas (http://www.proteinatlas.org)^75, 140^. **i**, Overlap between the top 100 human endothelial and mural cell subtype markers and those identified in mice. A more lenient set of 500 (instead of 100) mouse markers^37^ were used for comparison to ensure claims of species-specificity were robust. Note: the species-conservation of a cell type marker depends on species-specific changes in the given cell type *and* changes amongst the remaining background cell types.

We next ordered and aligned endothelial nuclei along a single one-dimensional Monocle pseudotime^74^ range to better recapitulate the anatomical arteriovenous axis. As expected, known arterial and venous markers peaked at opposite ends of this range, and capillary markers peaked in between (Fig. 2b). We used the 665 most significantly variable cluster genes to order the endothelial nuclei and observed a distribution of seven gradually changing gene expression patterns, representing arterial, capillary, or venous segments, and combinations thereof (Fig. 2c). We confirmed that the Monocle range represented a cell order matching anatomical arteriovenous zonation by examining data from the Human Protein Atlas^75^ (Supplemental Fig. 9a). Our patterns recapitulate the gradual/ seamless zonation continuum described in mice—but interestingly, this similar overall continuum arises from significantly different individual/ component zonation markers (Supplemental Table 2, 5). For example, only a minority of the top 100 human arterial, capillary, and venous markers are such in mice (Fig. 2j), even if we expand the denominator of mouse genes compared against to 500 genes.

We thus wondered whether established zonation markers in mice would be conserved in humans. We calculated a score (Methods) measuring each gene’s specificity to a given zonation (e.g., arterial, capillary, venous). Indeed, we observed across all vessel segments a significant number of markers that lost their predictive value between species (Fig. 2d, Supplemental Fig. 10). For example, the blood clotting gene von Willebrand factor (*VWF*) is largely expressed in mouse venous endothelial cells^37^. However, in humans, *VWF* is highly expressed throughout the endothelium, even in small diameter capillaries (Fig. 2d-e). VWF abundance has been tied to increased risk for ischemic stroke^76, 77^, and its species-specific distribution could be one reason why mouse models of stroke have faced notoriously low translational success rates^78, 79^.

We next visualized and clustered 34,508 mural cell nuclei in UMAP space, resolving clusters for arterial smooth muscle cells (aSMCs) and arteriolar SMCs (aSMCs)—but interestingly, we also discovered two subclusters of pericytes (Fig. 2f, Supplemental Fig. 8c-e). One pericyte subcluster was enriched for small molecule transmembrane transport activity, which we refer to as T-pericytes (for transport); while the other for extracellular matrix (ECM) formation and regulation, which we refer to as M-pericyte (for matrix). This suggests that function rather than anatomical location is the major driver of pericyte transcriptional identity in humans, and that both capillary and venous (vSMC) pericytes span these two functional clusters (confirmed below, Fig. 2g-h, Supplemental Fig. 9b, d). The existence of an M-pericyte cluster holds interesting implications for small vessel diseases like CADASIL, CARASIL, and Collagen IV deficiencies for which perturbations in the vascular ECM cause disease^80, 81^. Moreover, human T-pericytes—but not mouse pericytes—express transporters like the GABA transporter 1 *SLC6A1* (involved in epilepsy) and glutamate transporter *SLC1A3* (Supplemental Fig. 9c), suggesting an evolutionary pressure for expanded solute transport across the human BBB. Because recent mouse pericyte datasets have reported confounding endothelial contamination^37, 71, 82, 83^, we assessed and found no such contamination in our human pericyte nuclei (Supplemental Fig. 3, 7b). The isolation of nuclei instead of whole cells may aid in minimizing contaminating endothelial fragments.

To study mural cell zonation, we similarly compared the distribution of known mural cell transcripts across the Monocle range with corresponding protein expression *in situ* (Fig. 2g, Supplemental Fig. 9b, d). We used the 799 most significantly variable cluster genes to order all 36,825 mural cell nuclei and observed the expected order of aSMC markers on one end (e.g., *ACTA2*, *TAGLN*), followed by aaSMC (e.g., *CTNNA3*, *SLIT3*); and pericyte markers on the other end (e.g., *ABCC9*, *PTN*). Recapitulating the mural cell pattern described in mice^37^—and as opposed to the gradual zonation pattern in endothelial cells—, we observe in human mural cells an abrupt transition between SMCs and pericytes: one set of transcripts are expressed highly in aSMCs and aaSMCs but at low levels in pericytes, while another set of transcripts exhibits the opposite pattern (Fig. 2g). As expected from their clustering by functional pathways, the two pericyte subclusters did not segregate along the Monocle range and localized across both large and small diameter vessels *in situ*, suggesting that they intercalate throughout the capillary and venous vasculature (Fig. 2h). Moreover, previously reported vSMC markers^37^ were expressed in both pericyte clusters (Supplemental Fig. 9d).

Given species-specific differences across all brain cell types, we find that only a minority of the top mouse SMC and pericyte markers retain their predictive value in humans (Fig 2j, Supplemental Fig. 9d). Because the proteins encoded by zonation marker genes perform a variety of important functions at defined arteriovenous locations, species-specific endothelial and mural cell differences likely reflect fundamental differences in brain vascular properties that can now be tested to inform translational studies.

### Molecular definitions for perivascular and meningeal fibroblasts

Complex barrier structures maintain brain homeostasis^84^. Cooperating with the vascular BBB, the recently (re)discovered meningeal lymphatics plays important roles in waste clearance and neuroimmune surveillance^85–88^. Using annotations from recent mouse studies^42, 89, 90^, we noticed our capture of fibroblast-like cells from both the meningeal and vascular barriers provided the opportunity to directly compare these populations for insights into their specialized functions (Fig. 3a-e). First, we noticed that fibroblasts transcriptionally segregated according to anatomical location: vascular versus meningeal (Fig. 3a-b), but also separated according to the layers of the meninges (Fig. 3a, d). No strong differences were observed between hippocampal and cortical-derived fibroblasts (Supplemental Fig. 11b), suggesting that the micro- but not macro-environment shapes brain fibroblast identity.

**Figure 3.**
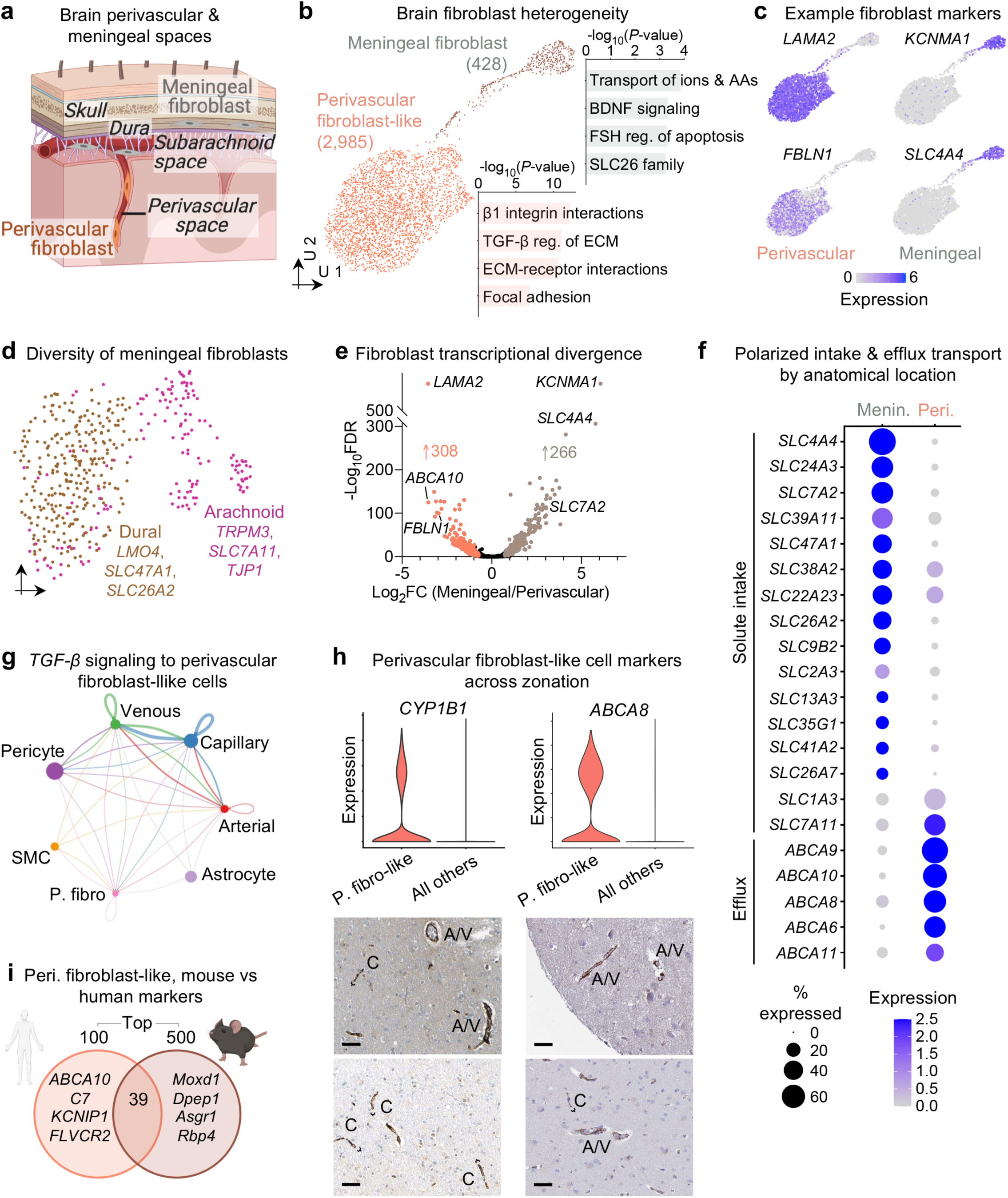
**Molecular definitions for brain perivascular and meningeal fibroblasts.** **a**, Anatomical reference of the human meninges (dura and arachnoid) and perivascular space, each with a resident fibroblast population. **b**, UMAP of 2,985 human perivascular fibroblast-like nuclei and 428 meningeal fibroblast nuclei. Enriched biological pathways derived from respective fibroblast cell type markers (Supplemental Table 2). **c**, Expression of example markers demarcating perivascular from meningeal fibroblasts. **d**, UMAP of 428 meningeal fibroblast nuclei, subclustering into anatomically segregated dural and arachnoid space fibroblasts. **e**, Differentially expressed genes (DEG) comparing perivascular and meningeal fibroblasts (MAST, Benjamini Hochberg correction; FDR < 0.01 and logFC>0.5 [log_2_FC>0.72] to be colored significant). **f**, Expression of all differentially expressed (from (**d**)) SLC and ABC family members across perivascular and meningeal fibroblasts. **g**, Circle plot showing the number of statistically significant intercellular signaling interactions among cerebrovascular cells for the *TGF-β* family of molecules (permutation test, CellChat^93^). **h**, Expression of perivascular fibroblast-like cell-specific markers *CYP1B1* and *ABCA8* compared to all other CNS cell types (top). Immunohistochemical co-localization (bottom) of CYP1B1 and ABCA8 to small diameter (< 10 microns, black brackets, ‘C’) capillaries in the human brain. Scale bar = 50 microns. **i**, Overlap between the top 100 perivascular fibroblast-like cell markers and those identified in mice. A more lenient set of 500 (instead of 100) mouse markers^37^ were used for comparison to ensure claims of species-specificity were robust. Note: the species-conservation of a cell type marker depends on species-specific changes in the given cell type *and* changes amongst the remaining background cell types.

Pathway enrichment analysis of marker genes demonstrated a strong divergence in fibroblast functions by anatomical location (Fig. 3b, e): perivascular fibroblast-like cells showed enriched expression for ECM structural components or its modifiers and receptors (e.g., “TGF- β regulation of the ECM), while meningeal fibroblasts enriched for solute transporters. This suggests that perivascular but not meningeal fibroblasts form fibrotic scars after brain injury^91, 92^. Closer comparison of differentially expressed genes between fibroblast populations (Fig. 3e) revealed a remarkable polarization of solute influx and efflux pumps: meningeal fibroblasts specifically expressed SLC influx solute transporters, while perivascular fibroblasts exclusively expressed ABC efflux pumps (Fig. 3f). Perivascular fibroblasts reside in the Virchow-Robin space, and thus like meningeal fibroblasts, come into contact with the cerebrospinal fluid (CSF). This cooperative circuit of polarized transporters suggests fibroblast regulation of solute exchange between the brain and CSF. A gradient of polarized influx/ efflux within a shared CSF compartment also provides evidence for convective rather than diffusive fluid flow via the recently described ‘glymphatic’ system^85^.

Because perivascular fibroblast-like cells reside in close proximity to other vascular cells captured, we used our single-cell data to infer cell-cell communication pathways^93, 94^. This corroborated fibroblast-like cells as major recipients of TGF-β signaling (Supplemental Fig. 11c-d). Overstimulation of TGF-β signaling in the brain vasculature promotes ECM basement membrane thickening and triggers neuropathology, though the effector cell type has been unknown^95, 96^. This analysis nominates perivascular fibroblast-like cells. Cell-cell communication analysis also predicted signaling between capillaries and fibroblast-like cells, despite their localization in mice exclusively around arterial and venous vasculature^37^. Using fibroblast-specific genes conserved in mice and humans, we surprisingly found fibroblast-like cell marker co-localization around human capillaries (<10 μM diameter) as well as larger vessels *in situ* (Fig. 3h, Supplemental Fig. 12). Fibroblast or fibroblast-derived protein localization in capillaries poses interesting questions for vessel development and maintenance. As in endothelial and mural cells, perivascular fibroblast-like cell markers varied by species (Fig. 3j). Together, these data provide a first characterization of human brain fibroblast diversity, revealing the molecular basis of their anatomical specialization and a cooperative circuit for CSF solute exchange.

### Vascular cell-type specific perturbations in Alzheimer’s disease

Alzheimer’s disease (AD) is a progressive neurodegenerative disorder culminating in severe impairment of memory, cognition, and executive functions^97, 98^. These impairments arise from complex perturbations in cell composition and gene expression^99–101^. We thus sought to profile changes in the AD human brain vasculature at single-cell resolution. We defined our AD patient group by clinical diagnosis and confirmed via immunohistochemistry the presence of β-amyloid plaques in the hippocampus and cortex (Supplemental Fig. 13).

Recent studies have identified context-dependent, disease associated glial subpopulations^28–30, 67, 102^. We did not observe new vascular cell subclusters emerging with AD (Fig. 4a, Supplemental Fig. 8). However, in contrast to parenchymal cells^28, 29^, we found a strong loss of brain vascular nuclei—across endothelial, SMC, pericyte, and fibroblast-like cells—with AD (Fig. 4a-b). This is consistent with reports of focal cerebrovascular damage in AD^27, 103^, but suggests that vascular loss is widespread across cells types throughout the arteriovenous axis. Interestingly, among pericytes, only M-pericytes involved in ECM organization declined (Fig. 4b). This selective, disease-associated susceptibility provides a molecular hypothesis for the physical BBB breakdown and vascular ECM disruption reported in AD^104^. Finally, because the observed reductions in vascular nuclei could also reflect an increased fragility during isolation, we stained for and confirmed our findings *in situ* (Fig. 4c). In short, instead of a new AD-associated subpopulation emerging, we find in the vasculature an AD-associated disappearance of specific cell type subpopulations.

**Figure 4.**
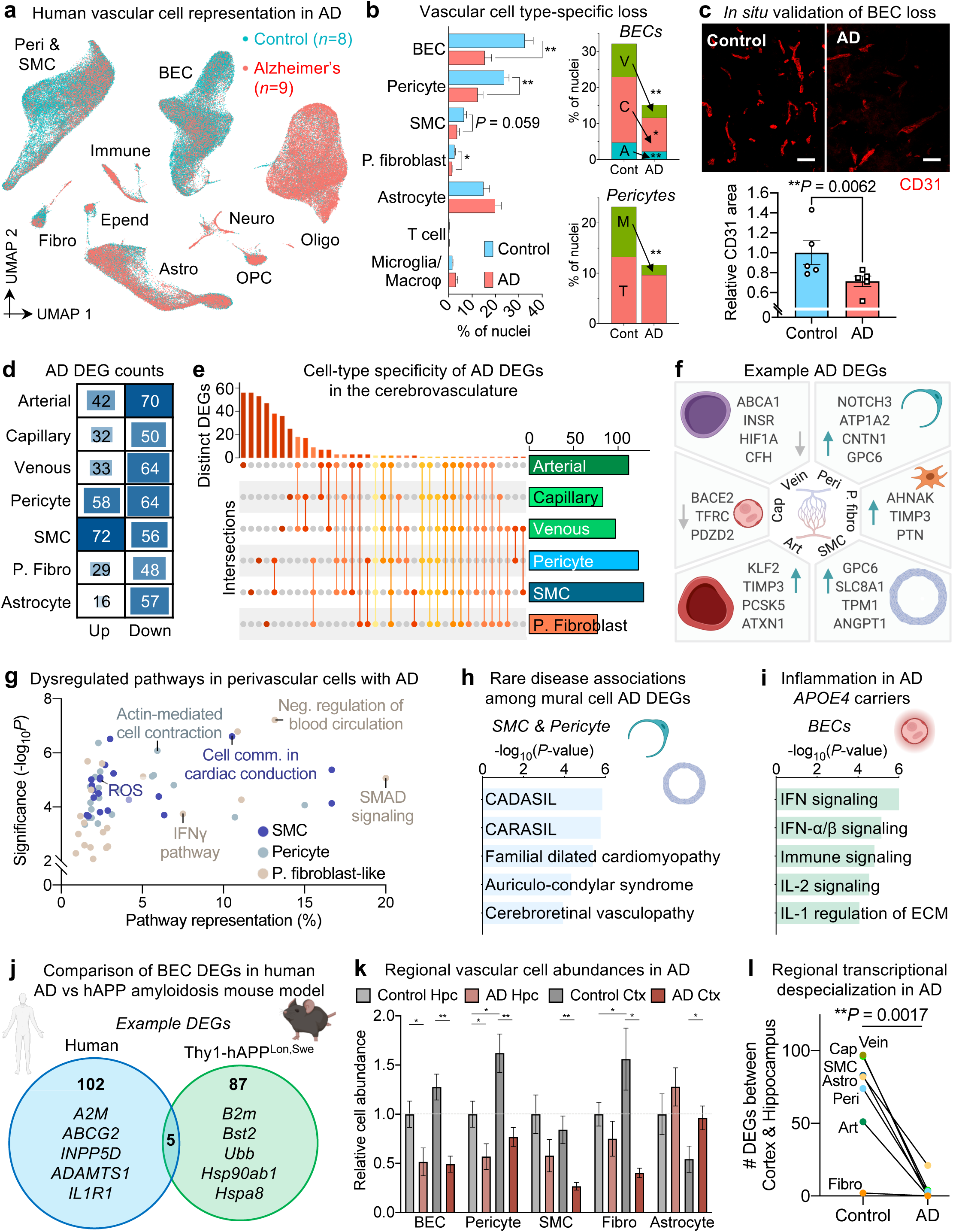
**Vascular cell-type specific perturbations in Alzheimer’s disease.** **a**, UMAP of 143,793 nuclei captured from 17 human hippocampus and superior frontal cortex samples, colored by Alzheimer’s disease (AD) diagnosis. **b**, Proportion of cell types captured in AD and controls (left). Proportion of brain endothelial vessel segments and pericyte subpopulations in AD and controls (right) (*n* = 8 controls, *n* = 9 AD, two-sided t-test; mean +/- s.e.m.). **c**, Immunohistochemical validation of a loss of BECs in AD. Scale bar = 50 microns (*n* = 5 controls and AD, nested two-sided t-test; mean +/- s.e.m.). **d**, Differentially expressed gene (DEG) counts for each cell type in AD. The intensity of the blue color and the size of the squares are proportional to entry values. **e**, Matrix layout for intersections of AD DEGs shared across and specific to each cell type. Circles in the matrix indicate sets that are part of the intersection, showing that most DEGs are cell type-specific. **f**, Example differentially expressed genes (DEGs) in AD: arterial (Art), capillary (Cap), venous (Vein), pericyte (Peri), perivascular fibro blast-like cell (P. fibro), and smooth muscle cell (SMC). Blue arrow indicates upregulated and grey arrow downregulated genes. **g**, Enriched biological pathways from AD differentially expressed genes in pericytes, smooth muscle cells, and perivascular fibroblast-like cells, plotted by Pathway Representation (in a given pathway, what proportion of all members are DEGs) and Significance (-log_10_*P*) of pathway enrichment. **h**, Rare diseases associated with differentially expressed genes in pericytes and smooth muscle cells. **i**, Enriched biological pathways from genes upregulated in AD *APOE4* carriers in capillary and venous endothelial cells. **j**, Venn diagram comparing DEG BECs in human AD samples compared to those from the Thy1-hAPP T41B^Lon,Swe^ amyloidosis mouse model^112^. Note that only genes with human-mouse orthologs are shown, and that the absolute logFC threshold for calling DEGs in mouse APP BECs was lowered to 0.15 (by half) to ensure claims of limited overlap with human BECs were robust. **k**, Among patients with both hippocampus and superior frontal cortex profiled (*n*=4 controls and *n*=4 AD), quantification of the relative abundance of major vascular cell types (control hippocampus set as reference, unpaired two-sided t-test; mean +/- s.e.m.). **l**, As in **(k**), but comparison of the number of DEGs between brain regions for each cerebrovascular cell type. Analysis done separately for control and AD samples (*n*=7 cell types, unpaired two-sided t-test; mean +/- s.e.m.)

We next systematically examined cell type-specific gene expression changes in AD (Methods). Across the vascular cell types captured in sufficient numbers for statistical power, we identified 463 unique differentially expressed genes using more stringent thresholds (DEGs, Methods, Fig. 4d, Supplemental Table 6). Overall, mural cells exhibited the strongest changes, with other cell types showing a signature of gene repression: 61-78% of DEGs were downregulated (Fig. 4d). DEGs were robustly detected across different levels of expression (Supplemental Table 6). The vast majority of DEGs were cell type- (Fig. 4e-f) and zonation-specific, suggesting a heterogenous response to AD pathology across the vasculature. Intriguingly, several DEGs are risk genes implicated in recent AD and small vessel disease GWAS studies (Fig. 4f). At the pathway level, DEGs in mural cells and fibroblast-like cells implicated dysregulated vasoconstriction and compromised blood flow (Fig. 4g). This provides a molecular basis for the cerebral hypoperfusion discernible in MRI-based imaging of living AD patients^105, 106^. Interestingly, DEGs in pericytes and SMCs resembled those found in CADASIL and CARASIL^107–110^ (Fig. 4h), rare hereditary diseases also marked by impaired blood flow, cognitive deficits, repeated strokes, and dementia.

APOE4 carriers have been reported to exhibit accelerated BBB breakdown before cognitive impairment^11, 111^, though the underlying mechanisms are unclear in humans. With patient *APOE* genotypes, we performed similar DEG analysis, finding dramatic interferon inflammation in the endothelium of *APOE4* carriers (Fig. 4i, Supplemental Fig. 14, Supplemental Table 7). Finally, we sought to understand the overlap between human vascular AD DEGs and those in mouse models of AD. Such models have facilitated mechanistic study of β-amyloid pathology, but recent reports describe significant species differences in various cell types, such as microglia^30, 44, 102^. We isolated brain endothelial cells from 12-14 month old Thy1-hAPP^Lon,Swe^ mice (and littermate wild-type controls)^112^ that present prominent amyloidosis, neuroinflammation, and synapto-dendritic degeneration—and processed them for single-cell sequencing. Surprisingly, we observed minimal overlap between human AD and mouse hAPP DEGs in brain endothelial cells (Fig. 4j).

AD pathology begins and spreads via a strikingly consistent regional pattern^98, 113^. We thus assessed the impact of AD on brain regional vascular specialization (Fig. 1e). Within controls, we found greater vascular density in the cortex compared to hippocampus (Fig. 4k), reflecting either regional baseline differences or hippocampal deficits with normal aging^114^. We found these regional differences erased in AD patients (Fig. 4k). Likewise, by comparing the number of DEGs between the cortex and hippocampus of the same patients, we noticed a global loss of brain regional specialization across vascular cell types in AD patients (Fig. 4l)— suggesting impairments in brain region-specific vascular function. Together, these findings show that AD patients exhibit heterogeneous cell type-, zonation-, region-, and species-specific perturbations across the brain vasculature that require dedicated isolation and single-cell approaches to profile.

### AD GWAS disease variants enriched in the human brain vasculature

A major goal of biomedical research is to identify genes that cause or contribute to disease. GWAs studies have shed insight into the molecular pathways contributing to AD^115, 116^, though the cell type context in which GWAS genes are expressed was long unknown. Recent snRNA-seq and other cell type-resolved studies have strongly implicated microglia as the major AD GWAS-expressing cell type^28–30, 64, 116–121^. We wondered, however, whether the unintended depletion of brain vascular cells in conventional preparations may have prematurely dismissed such evidence in brain vascular cells. We curated recent AD GWAS studies^117–119, 122, 123^ to identify and order the top 45 risk genes. With our more comprehensive survey of brain cell types, we calculated the cell type proportional expression for each GWAS gene using Expression Weighted Cell Type Enrichment (EWCE)^124^. We indeed observed among brain parenchymal cells a specific myeloid signature for top AD GWAS genes such as *TREM2*, *MS4A6A*, *CR1*, and *SPI1* that are now the subject of intense mechanistic study (Fig. 5a, right).

**Figure 5.**
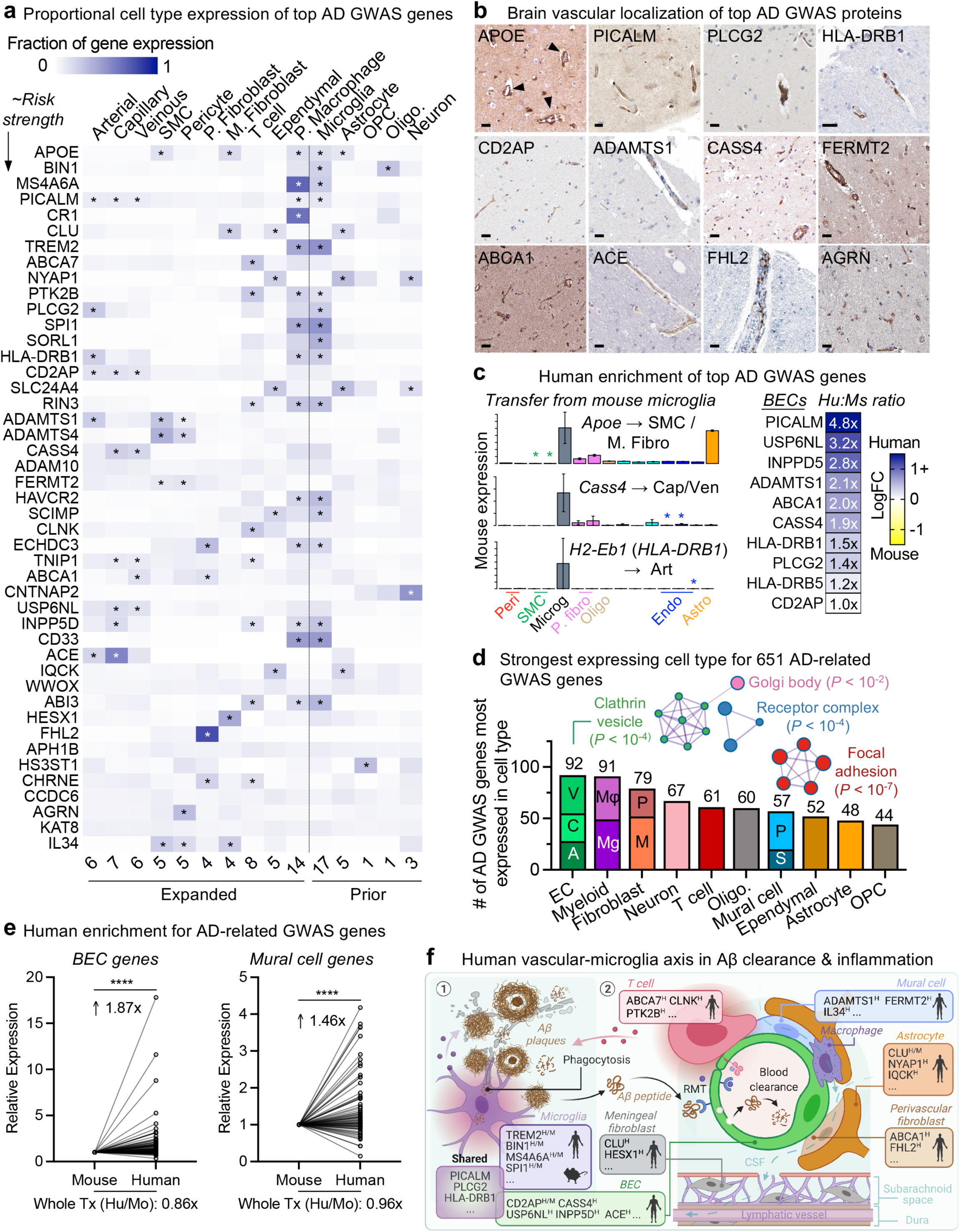
**GWAS disease variants are enriched in the human brain vasculature.** **a**, Proportional expression of the top 45 AD GWAS genes across all major brain cell types. Expression values for a given gene sums to 1 across cell types using the EWCE method^124^. Genes ordered in approximate risk strength^117–119, 123^. Asterisks denote strongest expressing cell types. Cells to the left of dashed line are from the vasculature, newly added here; to the right, parenchymal cells captured before. Numbers on the bottom summarize the number of GWAS genes enriched in a given cell type. Note: *MSA46A* represents the average expression of *MS4A46A*, *MS4A4A*, and *MS4A4E*; likewise, *HLA-DRB1* averages *HLA-DRB1* and *HLA-DRB5*. As in prior human brain RNA-seq datasets^60^, *EPHA1* was not robustly detected. **b**, Immunohistochemical confirmation of vascular localization of proteins encoded by 12 top AD GWAS genes from (**a**). Scale bar = 25 microns. Arrowheads in APOE point to signal around larger diameter vessels, consistent with predicted SMC expression. Images from the Human Protein Atlas (http://www.proteinatlas.org)^75, 140^. **c**, Enrichment of top AD GWAS genes in human over mouse endothelial cells (left). BEC heatmap of top AD GWAS genes is colored by logFC(human/mouse) and labeled by the linear fold-change (human/mouse) value. Example genes highly expressed in or specific to microglia in mice that are then expressed in vascular cells in humans (right). **d**, Quantification of the number of AD and AD-related trait GWAS genes^29^ most expressed in a given cell type. 383 of 651 genes (59%) mapped to vascular or perivascular cell types. PPI network of GO Cellular Components enriched in BECs at significance *P* < 0.05 (Metascape^141^). A = arterial, C = capillary, V = venous endothelial cell (EC). Mg = microglia, and Mφ = macrophage. In fibroblasts, M = meningeal and P = perivascular. In mural cells, S = SMC and P = pericyte. **e,** Human enrichment of AD-related trait GWAS genes^29^ highest expressed in BECs (left) and mural cells (right). In contrast to GWAS genes, the ratio of human to mouse expression across the whole transcriptome is less than or ∼1 for both cell types (bottom, paired two-sided t-test, \*\*\*\**P* < 0.0001 and \*\*\**P* = 0.0002). **f**, Summary of AD GWAS genes enriched in microglia and vascular cells mediating common pathways in protein clearance and inflammation. Mouse and human superscripts denote whether expression has been confirmed in that species for a given gene. Proposed model is described in Discussion.

Intriguingly, we noticed that several GWAS genes were strongly expressed in human brain vascular and perivascular cell types (Fig. 5a, left, Supplemental Fig. 15). This included the two GWAS genes previously implicated in the mouse vasculature, *PICALM* and *CD2AP*^64, 122^. But this also included other surprising genes, such as the immune-related *PLCG2* and *HLA-DRB1/5* in arterial cells, the endocytic *INPP5D* and *USP6NL* in capillaries, and ECM-related *ADAMTS1*, *ADAMTS4*, *FERMT2*, and *AGRN* in SMCs and pericytes (Fig. 5a). Within pericytes, expression varied across M- and T-pericyte subtypes (Supplemental Fig. 16a). *APOE*, often linked to myeloid cells and astrocytes, was robustly expressed in human SMCs and meningeal fibroblasts. Remarkably, several GWAS genes like *ABCA7* and *CLNK* were enriched in perivascular T cells. Consistent with our findings, an independent dataset shows minimal expression of these genes in parenchymal brain cell types (Supplemental Fig. 16b). Likewise, several GWAS genes like *ABCA1*, *FHL2*, *HESX1*, and *IL34* were enriched in perivascular and meningeal fibroblasts. Importantly, we confirmed our findings via immunohistochemical staining. We observed vascular localization for 12 proteins encoded by predicted-vascular GWAS genes, such as CASS4, FERMT2, ACE, PLCG2, and FHL2 (Fig. 5b). Most GWAS genes exhibited minor expression differences between the hippocampus and cortex (Supplemental Fig. 16c). In total, at least 30 of the top 45 AD GWAS genes are enriched in cells of the human brain vasculature (not including those solely in perivascular macrophages). Their distribution across all vascular cell types suggests that vascular and perivascular involvement in AD pathology may be more thorough and complex than anticipated.

The human brain vascular expression of putatively parenchymal—especially myeloid— AD GWAS genes made us wonder whether these genes are expressed in different cell types between mice and humans. We thus examined the expression of human vascular GWAS genes in mouse datasets^90^. Indeed, many genes like *APOE*, *CASS4*, *INPP5D*, and *HLA-DRB1* were predominately expressed in microglia in mice but then also exhibited vascular expression in humans (Fig. 5c, Supplemental Fig. 16d). Corroborating this, nearly every top GWAS gene expressed in BECs exhibited greater expression in humans than in mice (Fig. 5c). Together, these data suggest a partial evolutionary transfer of AD risk genes and pathways from microglia to the vasculature from mice to humans, with implications for translational studies.

We next broadened our scope to a previously compiled list of hundreds of GWAS genes for AD and AD-related traits (Supplemental Fig. 17a-b)^29^. We observed robust expression across vascular and perivascular cell types (Supplemental Fig. 17a-b). Using EWCE analysis, we found that many risk genes were expressed cell type-specifically (Supplemental Fig. 17). For each gene, we assigned the cell type with the strongest expression, discovering surprisingly, that endothelial cells harbored the most AD-related GWAS genes, followed by microglia/ macrophages (Fig. 5d, Supplemental Table 8). Within BECs, AD-related GWAS genes (Fig. 5a-b) enriched for protein endo- and transcytosis components, such as receptor and clathrin vesicle components (Fig. 5d). We recently demonstrated a decline in BEC clathrin-mediated transcytosis^15^ with age, suggesting one mechanism by which aging and GWAS genes converge to impair β-amyloid clearance and increase AD risk. In total, over half of all AD-related GWAS genes mapped to vascular or perivascular cell types (383 of 651).

As with top AD GWAS genes, we observed enhanced human over mouse expression of AD-related genes in both BECs and pericytes (Fig. 5e). Importantly, this human-enhanced expression is not observed for the whole transcriptome. Together, these data provide a more comprehensive understanding of the cell types contributing to AD risk. We suggest that a vascular-microglia axis underlies the genetic risk for AD via shared protein clearance (BEC-microglia) and inflammatory pathways (BEC-T cell-microglia) (Fig. 5f), and that this axis is evolutionarily expanded in humans.

## Discussion

We report here 143,793 single-cell, genome-wide quantitative transcriptomes from the human brain vasculature in health and AD. We use these transcriptomes to molecularly define the principal vascular cell types; their differences by brain region and species; the organizational principles of endothelial, mural, and fibroblast-like cells; a selective loss of M-pericytes and the transcriptomic perturbations contributing to clinical AD dementia; and the unexpected expression of AD GWAS genes across the human brain vasculature. We subsequently confirm these findings *in situ* at the protein level. Single-cell resolution was necessary for these findings, which would have been obscured in the average profiles generated via bulk RNA-sequencing.

How do human vascular GWAS genes fit into established AD pathways? Current understanding implicates β-amyloid metabolism, cholesterol/ lipid dysfunction, innate immunity, and endocytosis^121, 125^. Vascular GWAS gene expression confirms these pathways and expands the set of cell types involved, such as β-amyloid endocytosis and clearance via BEC clathrin-mediated transport; and adaptive in addition to innate immunity via perivascular T cells (Fig. 5f). We propose that the dramatic expansion of the human brain, brain activity, and activity byproducts (like β-amyloid^126^) necessitates enhanced neuroimmune surveillance and clearance mechanisms. In this model, microglia are still frontline participants in AD pathogenesis. But more so than in mice, human vascular and perivascular cells partake. For example, the clearance functions of microglia can become overwhelmed^127^, diverting debris clearance to BECs. This is supported by recent studies finding microglial depletion results in cerebral amyloid angiopathy^128^. But unlike microglia^129^, vascular cells are unable to proliferate efficiently^130^. Thus, constant vascular exposure to β-amyloid triggers dysfunction via cell loss and impaired blood and CSF flow^131^ (Fig. 4). Recent work identified a human-unique CD8 TEMRA (CD45RA^+^CD27^−^) population clonally expanded in AD CSF^132^. Thus, it is possible that perivascular/ meningeal T cell GWAS hits contribute to inflammatory pathology in ways not seen in mice. Together, we suggest an intertwined microglia-vascular axis expanded in humans, with vascular cells playing an auxiliary role via shared endocytosis and inflammatory pathways. We note though the likelihood of additional vascular contributions, as evidenced by SMC, pericyte, and both perivascular and meningeal fibroblast-enriched GWAS genes of unclear function.

Given the evolutionary divergence between mice and humans, these data expand by orders of magnitude the number of arteriovenous markers in the human brain that can now be used for vessel type identification, to assess the fidelity of *in vitro* human cell and organoid cultures, and to reliably deconvolute and enhance the utility of hundreds of publicly available bulk brain RNA-seq datasets (Supplemental Table 2). Mechanistic studies in mouse models can now be combined with this dataset to systematically pinpoint the cell types and genes mediating core vascular functions such as cerebral autoregulation and BBB permeability that are often seen perturbed in disease by clinical imaging^10^. Despite recent landmark studies on human brain parenchymal cells^116^, many AD and other disease risk variants remain unmapped. Because such variants enrich in gene expression-regulating enhancer regions^133^ that undergo accelerated evolution^134^, they may exert their influence through human brain vascular cells. With further optimization, this method should be compatible with ATAC-, ChIP-, and PLAC-seq assays^135–137^ to address this possibility.

Our work opens several translational opportunities. This dataset informs ongoing efforts to develop ‘brain shuttles’ and other modalities to better deliver therapeutics to treat human brain disorders^15, 21^. This method facilitates study of the brain vasculature across a variety of disease conditions, such as stroke, multiple sclerosis, and even COVID-19^67^. Together, the field now has a near complete catalog of cell types in the human brain, which can be integrated into ongoing efforts such as the Human Cell Atlas^138^. As with recent high-profile snRNA-seq studies of the brain parenchyma^28–32, 139^, it will now be important to distinguish which of the vascular transcriptional perturbations observed in disease are responsive versus driving, clarify their links to various clinical and pathologic traits, and dissect the exact mechanisms by which vascular-expressed AD GWAS genes confer greater disease risk. Overall, the VINE-seq method introduced here and the ensuing single-cell data (https://twc-stanford.shinyapps.io/human_bbb) provide a blueprint for finally studying the molecular makeup of the human brain vasculature, promising further discoveries in health, disease, and therapy.

## Supporting information

SI Table 1 Patient samples

SI Table 2 Cell type markers

SI Table 3 Brain region enriched genes

SI Table 4 Mouse vs human BEC and pericyte nuclei

SI Table 5 Cell subtype markers

SI Table 6 AD DEGs

SI Table 7 ApoE4 DEGs

SI Table 8 AD and AD-related GWAS genes

## Acknowledgments

We thank T. Iram, E. Tapp, N. Lu, M. Haney, O. Hahn, M.J. Estrada, and other members of the Wyss-Coray lab for feedback and support; Hansruedi Mathys, David A. Bennett, and participants in the CSHL BBB 2021 meeting for valuable advice; and H. Zhang and K. Dickey for laboratory management. This work was funded by the NOMIS Foundation (T.W.-C.), the National Institute on Aging (T32-AG0047126 to A.C.Y., 1RF1AG059694 to T.W.-C), Nan Fung Life Sciences (T.W.-C.), the Bertarelli Brain Rejuvenation Sequencing Cluster (an initiative of the Stanford Wu Tsai Neurosciences Institute), and the Stanford Alzheimer’s Disease Research Center (P30 AG066515). A.C.Y was supported by a Siebel Scholarship. F.K. and A.K. are a part of the CORSAAR study supported by the State of Saarland, the Saarland University, and the Rolf M. Schwiete Stiftung. This study was supported by the AHA-Allen Initiative in Brain Health and Cognitive Impairment: 19PABHI34580007. The statements in this work are solely the responsibility of the authors and do not necessarily represent the views of the American Heart Association (AHA) or the Paul G. Allen Frontiers Group.

## Author contributions

A.C.Y. and T.W.-C. conceptualized the study. A.C.Y. devised the isolation method. M.W.M. provided and A.C.Y. organized tissue samples. D.L.P. and A.C.Y. performed tissue dissociations. N.S., R.T.V., D.G., K.C., and A.C.Y. prepared libraries for sequencing. R.T.V., F.K., A.K., C.A.M., M.B.C., R.P., A.S., N.K., and A.C.Y. performed computational analysis. D.P.L, C.A.M., M.A., D.G., E.Y.W., J.L., and A.C.Y. performed immunohistochemical stains. P.M.L. developed the searchable web interface (Shiny app). C.A.M. and A.C.Y. drew diagrams. A.C.Y. wrote the manuscript with input from all authors. A.C.Y. and T.W.-C. supervised the study.

## Competing interests

T.W.-C. is a founder and scientific advisor of Alkahest Inc.

## Methods

### Isolation of vascular nuclei from frozen post-mortem brain tissue

Post-mortem fresh-frozen hippocampus and superior frontal cortex tissue were obtained from the Stanford/ VA/ NIA Aging Clinical Research Center (ACRC) with approval from local ethics committees and patient consent. Group characteristics are presented in Supplemental Table 1. Note, patients were grouped by *clinical* diagnosis, with two of the control patients exhibiting amyloid beta plaque staining in the hippocampus, though not to a sufficient degree for an expert pathologist to diagnose Alzheimer’s disease by histopathological criteria. Clinical instead of pathologic diagnosis was chosen because of potentially vascular contributions to AD independent of the well-known hallmarks of AD, β-amyloid and tau pathophysiology^7^. All procedures were carried out on ice in a 4°C cold room as rapidly as possible. 0.3 grams or more of brain tissue was thawed on ice for 5 minutes with 5 ml of nuclei buffer (NB): 1% BSA containing 0.2 U μl^−1^ RNase inhibitor (Takara, 2313A) and EDTA-free Protease Inhibitor Cocktail (Roche, 11873580001). Tissue was quickly minced and homogenized with 7 ml glass douncers (357424, Wheaton) until no visible chunks of debris remained. Similar to before^54^, homogenates were transferred into 50 ml tubes containing 35 ml of chilled 32% dextran (D8821, Sigma) in HBSS. Samples were vigorously mixed before centrifugation at 4,400g for 20 minutes with no brake. After centrifugation, samples separate into a top myelin layer, middle parenchymal layer, and vascular-enriched pellet. The myelin layer was aspirated, tips changed, and the parenchymal layer carefully removed without disturbing the pellet. Pellets were resuspended in 8 ml of 32% dextran, transferred to 15 ml falcon tubes, and centrifuged again. Vascular-enriched pellets were gently resuspended in 1ml of NB and added to pre-wetted 40 μm strainer sitting atop 50 ml falcon tubes. From here diverging from prior protocol, strainers were washed with 10 ml of cold 0.32 M sucrose in PBS and 90 ml of PBS until flow through the strainers was unimpeded to deplete contaminating parenchymal cells from trapped microvessels. At this step, retained microvessels turn white in color, indicating removal of circulating blood cells. Strainers were switched to new collection 50 ml falcon tubes. Various techniques were tested and optimized to extract vascular cells from the isolated microvessels (e.g., enzymatic digestion, TisssueRuptor, sonication, etc.), but nearly all resulted in loss of nuclei integrity or low nuclei complexity (<50 median genes/ nuclei). Eventually, adapting a method for the isolation of murine splenocytes proved successful: vascular fragments were mashed four times through the cell strainer using the plunger end of a 3 ml syringe, with intermittent elution via 10 ml of 0.32 sucrose and 40 ml of PBS. Liberated vascular cells were pelleted at 500g for 10 minutes and resuspended in 1.5 ml of EZ Prep Lysis Buffer (Sigma, NUC101) spiked with 0.2 U μl^−1^ RNase inhibitor (Takara, 2313A) and EDTA-free Protease Inhibitor Cocktail (Roche, 11873580001). Nuclei were homogenized with 2 ml glass douncers (D8938, Sigma) 20 times with pestle B (pestle A optional). Spiked EZ lysis buffer was added to samples up to 4 ml and incubated on ice for 5 minutes before pelleting at 500g for 6 minutes. This incubation step was repeated. Debris was depleted via a sucrose gradient before flow cytometry isolation of nuclei. Briefly, pelleted nuclei were resuspended in 0.5ml of NB before the addition of 0.9 ml of 2.2 M sucrose in PBS. This mixture was layered atop 0.5 ml of 2.2 M and samples were centrifuged at 14,000g for 45 minutes at 4°C, with no brake. Pellets were aspirated in 1ml of NB, filtered through a 40 μm strainer (Flowmi), transferred to FACS tubes, stained with Hoechst 3342 (1:2000, Thermo) and rabbit monoclonal anti-NeuN Alexa Fluor® 647 (1:500, Abcam, ab190565), and nuclei collected on a SH800S Cell Sorter into chilled tubes containing 1 ml of NB without protease inhibitor. In pilot runs, we noticed the cytometer overestimated nuclei counts by ∼3.4x, and thus we sorted ∼34,000 nuclei to target ∼10,000 nuclei per sample. Sorted samples were inspected for lack of debris on a brightfield microscope. We note that an iodixanol gradient^52^ can substitute for the 2.2 M sucrose, but that unfortunately with either gradient, flow sorting is required—unlike parenchymal myelin debris, vascular debris is not sufficiently removed by gradient centrifugation alone. Vascular debris will confound downstream cDNA traces with higher background and low molecular weight peaks. We are happy to share the detailed protocol widely but note that since high-quality human postmortem brain tissue is difficult to obtain, tissue would be limited in quantities to share widely.

### Droplet-based snRNA-sequencing

For droplet-based snRNA-seq, libraries were prepared using the Chromium Single Cell 3ʹ v3 according to the manufacturer’s protocol (10x Genomics), targeting 10,000 nuclei per sample after flow sorting (Sony SH800S Cell Sorter). 15 PCR cycles were applied to generate cDNA before 16 cycles for final library generation. Generated snRNA-seq libraries were sequenced on S4 lanes of a NovaSeq 6000 (150 cycles, Novogene).

### snRNA-seq quality control

Gene counts were obtained by aligning reads to the hg38 genome (refdata-gex-GRCh38-2020-A) using CellRanger software (v.4.0.0) (10x Genomics). To account for unspliced nuclear transcripts, reads mapping to pre-mRNA were counted. As previously published, a cut-off value of 200 unique molecular identifiers (UMIs) was used to select single nuclei for further analysis^28^. As initial reference, the entire dataset was projected onto two-dimensional space using Uniform Manifold Approximation and Projection (UMAP) on the top 30 principal components^142^. Three approaches were combined for strict quality control: (1) outliers with a high ratio of mitochondrial (>5%, <200 features) relative to endogenous RNAs and homotypic doublets (> 5000 features) were removed in Seurat^143^; (2) after scTransform normalization and integration, doublets and multiplets were filtered out using DoubletFinder^144^; and (3) after DoubletFinder, nuclei were manually inspected using known cell type-specific marker genes, with nuclei expressing more than one cell type-specific marker further filtered^144^. For example, BEC nuclei containing any reads for the following cell type markers were subsequently filtered: *MOBP*, *MBP*, *MOG*, *SLC38A11*, *LAMA2*, *PDGFRB*, *GFAP*, *SLC1A2*, and *AQP4*. We note that the vascular nuclei in prior human single cell datasets exhibit contamination with other cell type-specific gene markers, potentially confounding downstream analysis. After applying these filtering steps, the dataset contained 143,793 high-quality, single nuclei.

### Cell annotations & differential gene expression analysis

Seurat’s Integration function was used to align data with default settings. Genes were projected into principal component (PC) space using the principal component analysis (RunPCA). The first 30 dimensions were used as inputs into Seurat’s FindNeighbors, FindClusters (at 0.2 resolution) and RunUMAP functions. Briefly, a shared-nearest-neighbor graph was constructed based on the Euclidean distance metric in PC space, and cells were clustered using the Louvain method. RunUMAP functions with default settings was used to calculate 2-dimensional UMAP coordinates and search for distinct cell populations. Positive differential expression of each cluster against all other clusters (MAST) was used to identify marker genes for each cluster^145^. We annotated cell-types using previously published marker genes^28, 30, 32, 146^. For brain endothelial cells, zonation specificity scores for each gene were calculated separately for arterial, capillary, and venous segments as in the following example for a given gene in capillaries:

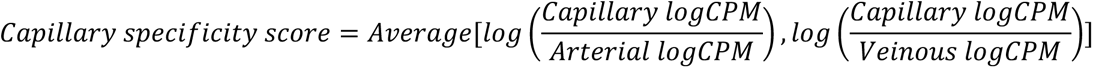

Differential gene expression of genes comparing Alzheimer’s disease, ApoE4, and control samples—or comparing cell type sub-cluster markers—was done using the MAST^145^ algorithm, which implements a two-part hurdle model. Seurat natural log (fold change) > 0.5 (absolute value), adjusted *P* value (Bonferroni correction) < 0.01, and expression in greater than 10% of cells in *both* comparison groups were required to consider a gene differentially expressed for subcluster analysis and natural log (fold change) > 0.3 (absolute value), adjusted *P* value (Bonferroni correction) < 0.01, and expression in greater than 10% of cells in both comparison groups for Alzheimer’s disease and ApoE4 comparisons, both more stringent than the default Seurat settings. We incorporated age, gender, and batch as covariates in our model. A more lenient threshold of the above but with natural log (fold change) > 0.2 (absolute value) was used for brain region (i.e., hippocampus vs cortex. Biological pathway and gene ontology enrichment analysis was performed using Enrichr^147^ or Metascape^141^ with input species set to *Homo sapiens*^141^. UpSet plots were generated using identified differentially expressed genes as inputs using the R package UpSetR^148^. Diagrams were created with BioRender.

### Monocle trajectory analysis

Monocle was used to generate the pseudotime trajectory analysis in brain endothelial and mural cells^74^. Cells were clustered in Seurat and cluster markers used as input into Monocle to infer arteriovenous relationships within endothelial cells and pericytes. Specifically, UMAP embeddings and cell sub-clusters generated from Seurat were converted to a *cell_data_set* object using SeuratWrappers (v.0.2.0) and then used as input to perform trajectory graph learning and pseudo-time measurement through Independent Component Analysis (ICA) with Monocle. Cluster marker genes identified in Seurat were used to generate a pseudotime route and plotted using the ‘plot_pseudotime_heatmap’ function.

### Cell-cell communication

Cell-cell interactions based on the expression of known ligand-receptor pairs in different cell types were inferred using CellChatDB^93^ (v.0.02). Briefly, we followed the official workflow and loaded the normalized counts into CellChat and applied the preprocessing functions *identifyOverExpressedGenes*, *identifyOverExpressedInteractions*, and *projectData* with standard parameters set. As database we selected the *Secreted Signaling* pathways and used the pre-compiled human Protein-Protein-Interactions as a priori network information. For the main analyses the core functions *computeCommunProb*, *computeCommunProbPathway*, and *aggregateNet* were applied using standard parameters and fixed randomization seeds. Finally, to determine the senders and receivers in the network the function *netAnalysis_signalingRole* was applied on the *netP* data slot.

### Mouse wild-type and APP T41B BEC single-cell and nuclei sequencing

Whole cell isolation from the CNS followed previously described methods^20, 40, 58^. Briefly, cortices and hippocampi were microdissected, minced, and digested using the Neural Dissociation Kit (Miltenyi). Suspensions were filtered through a 100 µm strainer and myelin removed by centrifugation in 0.9 M sucrose. The remaining myelin-depleted cell suspension was blocked for ten minutes with Fc preblock (CD16/ CD32, BD 553141) on ice and stained for 20 minutes with antibodies to distinguish brain endothelial cells (CD31^+^/ CD45^-^). Brain endothelial cells from 12-14 month old Thy1-hAPP^Lon,Swe^ mice and littermate wild-type control^112^ mice (pool of 4-6 mice per group) were sorted into PBS with 0.1% BSA. Nuclei isolation from 4-6 month-old mouse hippocampi followed protocols adapted from previous studies^28–30, 52, 149^. Briefly, tissue was homogenized using a glass douncer in 2 ml of ice-cold EZ PREP buffer (Sigma, N3408) and incubated on ice for 5 min. Centrifuged nuclei were resuspended in 1% BSA in PBS with 0.2 U μl^−1^ RNase inhibitor and filtered through a 40 μm cell strainer. Cells or nuclei were immediately counted using a Neubauer haemocytometer and loaded on a Chromium Single-Cell Instrument (10x Genomics, Pleasanton, CA, USA) to generate single-cell GEMs. The 10x-Genomics v3 libraries were prepared as per the manufacturer’s instructions. Libraries were sequenced on an Illumina NextSeq 550 (paired-end; read 1: 28 cycles; i7 index: 8 cycles, i5 index: 0 cycles; read 2: 91 cycles). De-multiplexing was performed using the Cellranger toolkit (v3.0.0) “cellranger mkfastq” command and the “cellranger count” command for alignment to the mouse transcriptome, cell barcode partitioning, collapsing unique-molecular identifier (UMI) to transcripts, and gene-level quantification. ∼70% sequencing saturation (>20,000 reads per cell) was achieved, for a median of ∼2,000 genes detected per cell and ∼16,500 genes detected in total. Downstream analysis using the Seurat package (v3)^150^ was performed as previously described^37^, applying standard algorithms for cell filtration, feature selection, and dimensionality reduction. Samples with fewer than 1,000 and more than 4,000 unique feature counts, samples with more than 15% mitochondrial RNA, samples with more than 15% small subunit ribosomal genes (Rps), and counts of more than 10,000 were excluded from the analysis. Genes were projected into principal component (PC) space using the principal component analysis (RunPCA). The first 30 dimensions were used as inputs into Seurat’s FindNeighbors and RunTsne functions. Briefly, a shared-nearest-neighbor graph was constructed based on the Euclidean distance metric in PC space, and cells were clustered using the Louvain method. RunTsne functions with default settings was used to calculate 2-dimensional tSNE coordinates and search for distinct cell populations. Cells and clusters were then visualized using 3-D t-distributed Stochastic Neighbor embedding on the same distance metric. Differential gene expression analysis was done by applying the Model-based Analysis of Single-cell Transcriptomics (MAST). Significant differentially expressed genes in Thy1-hAPP^Lon,Swe^ BECs were called by Log (fold change) > 0.15 (absolute value), adjusted *P* value (Bonferroni correction) < 0.01. This lowered Log (fold change) was to ensure our claims of limited overlap with human AD BECs were robust.

### *GWAS* analysis

For calculation of proportional cell type-specific gene expression, we followed the expression weighted cell type enrichment (EWCE) method described by Skene et al.^124^, and used previously on human snRNA-seq data^29^. For Alzheimer’s disease (AD) analysis, we compiled a list of top GWAS risk genes from Lambert et al.^117^, Kunkle et al.^118^, and Jansen et al.^119^, sorted descending by approximate *P*-value. Each gene’s expression sums to 1 across the cell types, with each heatmap cell showing the fraction of total gene expression as determined from EWCE analysis. The set of 720 AD and AD-related trait GWAS genes were obtained from Grubman, et al.^29^, and using EWCE analysis, the strongest expressing cell type was determined for each gene. Note that the original list was slightly parsed to 720, as several genes were not detected as expressed in our dataset.

For analysis across CNS diseases, from the GWAS catalog^151^, we obtained GWAS risk genes for neurological disorders [Alzheimer’s disease (AD), amyotrophic lateral sclerosis (ALS), brain aging, multiple system atrophy (MSA), multiple sclerosis (MS), Parkinson’s disease (PD), and narcolepsy], psychiatric disorders [Attention deficit hyperactivity disorder (ADHD), autism, bipolar disorder, depression, psychosis, post-traumatic stress disorder (PTSD), and schizophrenia], and neurobehavior traits [Anxiety, suicidality, insomnia, neuroticism, risk behavior, intelligence, and cognitive function]. We removed gene duplicates and GWAS loci either not reported or in intergenic regions and used a *P* < 9 × 10^−6^ to identify significant associations^29^. Then, since GWAS signals can point to multiple candidate genes within the same locus, we focused on the ‘Reported Gene(s)’ (genes reported as associated by the authors of each GWAS study). Following gene symbol extraction, we curated the gene set by (1) removing unknown or outdated gene names using the HGNChelper package (v.0.8.6), (2) converting remaining Ensembl gene IDs to actual gene names using the packages ensembldb (v.2.10.0) and EnsDb.Hsapiens.v86 (v.2.99.0), and (3) removing any remaining duplicates. For each disease, we allocated each of its GWAS risk genes to the cell type that proportionally expressed it most (EWCE analysis), before tallying this number in both counts and as a percentage of the diseases total number of GWAS risk genes. Finally, a statistical enrichment of each overlap against background was calculated using a hypergeometric test with the total background size set equal to the number of unique genes (21,306).

### Immunohistochemistry

Fresh-frozen control and AD human brain tissue (hippocampus and superior frontal cortex) adjacent to tissue processed for snRNA-seq was subjected to immunohistochemistry (IHC). 10 µm sections mounted on SuperFrost Plus glass slides were fixed with 4% paraformaldehyde (Electron Microscopy Services, 15714S) diluted in PBS at 4°C for 15 minutes before dehydration via an ethanol series or air drying. Sections were blocked in TBS++ (TBS + 3% donkey serum (130787, Jackson ImmunoResearch) + 0.25% Triton X-100 (T8787, Sigma-Aldrich)) for 1.5 hours at room temperature. Sections were incubated with primary antibodies at 4°C overnight: mouse monoclonal anti-CD31 (1:100, JC70A, Dako), rabbit polyclonal anti-VWF (1:100, GA527, Dako), rabbit polyclonal anti-SLC39A10 (1:100, HPA066087, Atlas Antibodies), rabbit polyclonal anti-ALPL (1:100, HPA007105, Atlas Antibodies), rabbit polyclonal anti-A2M (1:100, HPA002265, Atlas Antibodies), rabbit monoclonal anti-β-Amyloid (1:500, clone D54D2 XP, CST), and mouse monoclonal anti-Actin, α-Smooth Muscle - Cy3 (1:100, clone 1A4, Sigma). Sections were washed, stained with Alexa Fluor-conjugated secondary antibodies (1:250) and Hoechst 33342 (1:2000, H3570, Thermo), mounted and coverslipped with ProLong Gold (Life Technologies) or VECTASHIELD (Vector Laboratories before imaging on a confocal laser scanning microscope (Zeiss LSM880). Age-related autofluorescence was quenched prior to mounting with Sudan Black B, as before^15, 58^. National Institutes of Health ImageJ software was used to quantify the percentage of vasculature (CD31) or the predicted DEG SLC39A10 among CD31+ vasculature, following previously described protocols^15, 152, 153^. All analyses were performed by a blinded observer.

## Data Availability

Raw sequencing data is deposited under NCBI GEO: GSE163577. Data is also available to explore via an interactive web browser: https://twc-stanford.shinyapps.io/human_bbb.

## Supplemental Figures

**Supplemental Figure 1.**
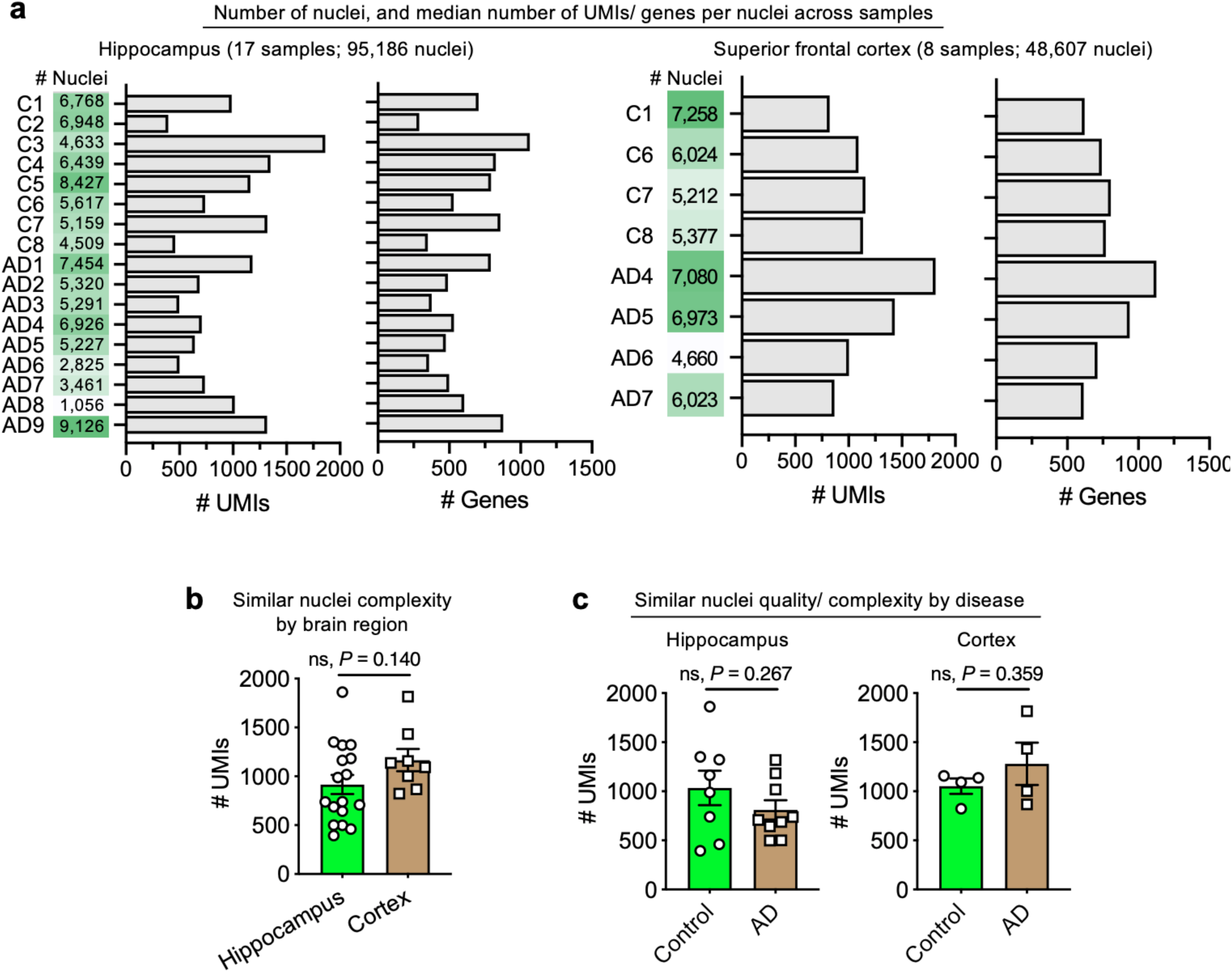
**Characterization of vascular nuclei captured.** **a**, Total number of nuclei, median number of unique molecular identifiers (UMI), and median number of genes for each human sample sequenced from hippocampus and superior frontal cortex. **b**, Quantification of the median number of genes detected per nuclei in medial frontal cortex and choroid plexus (*n*=17 hippocampus and *n*=8 cortex, two-sided t-test; mean +/- s.e.m.). **c**, Quantification of the median number of genes detected per nuclei in controls and Alzheimer’s disease (AD) samples in hippocampus and superior frontal cortex (*n*=8 controls and *n*=9 AD, two-sided t-test; mean +/- s.e.m.).

**Supplemental Figure 2.**
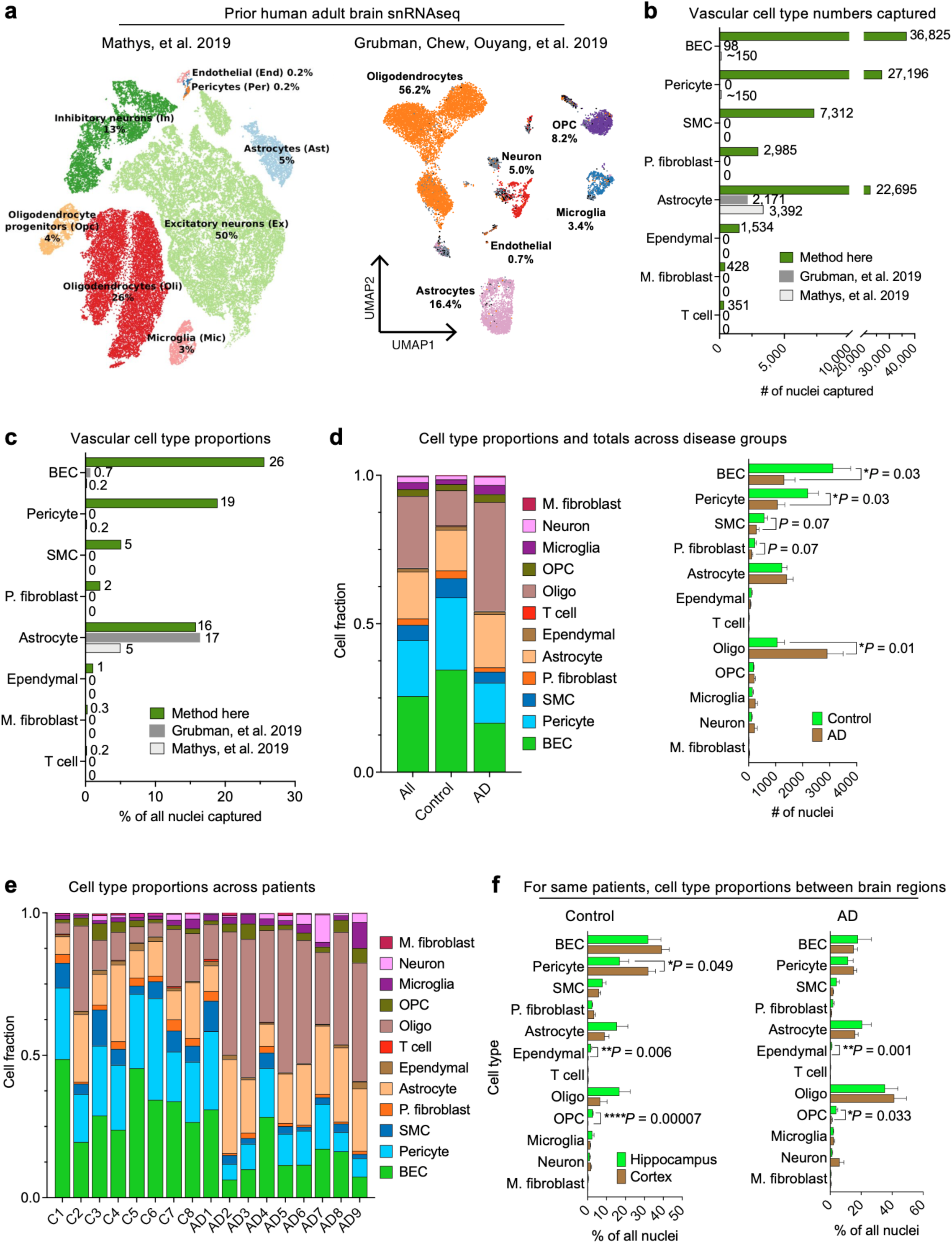
**Enhanced representation of brain vascular cells.** **a**, tSNE and UMAP projections of recent snRNA-seq studies^28, 29^ on post-mortem AD samples, capturing a relatively low representation of cerebrovascular cell types. **b**, **c**, Quantification of the number (**b**) and proportion (**c**) of cerebrovascular cell types captured via the method introduced here compared to recent snRNA-seq studies^28, 29^. **d**, Quantification of the proportion (left) and number (right) of captured cell types by control and AD patients (*n*=8 controls and *n*=9 AD, two-sided t-test; mean +/- s.e.m.). **e**, Quantification of the proportion of captured cell types per patient. **f**, For patients with both hippocampus and superior frontal cortex profiled, quantification of the proportion of cerebrovascular cell types captured in each region (*n*=4 controls and *n*=4 AD, two-sided t-test; mean +/- s.e.m.).

**Supplemental Figure 3.**
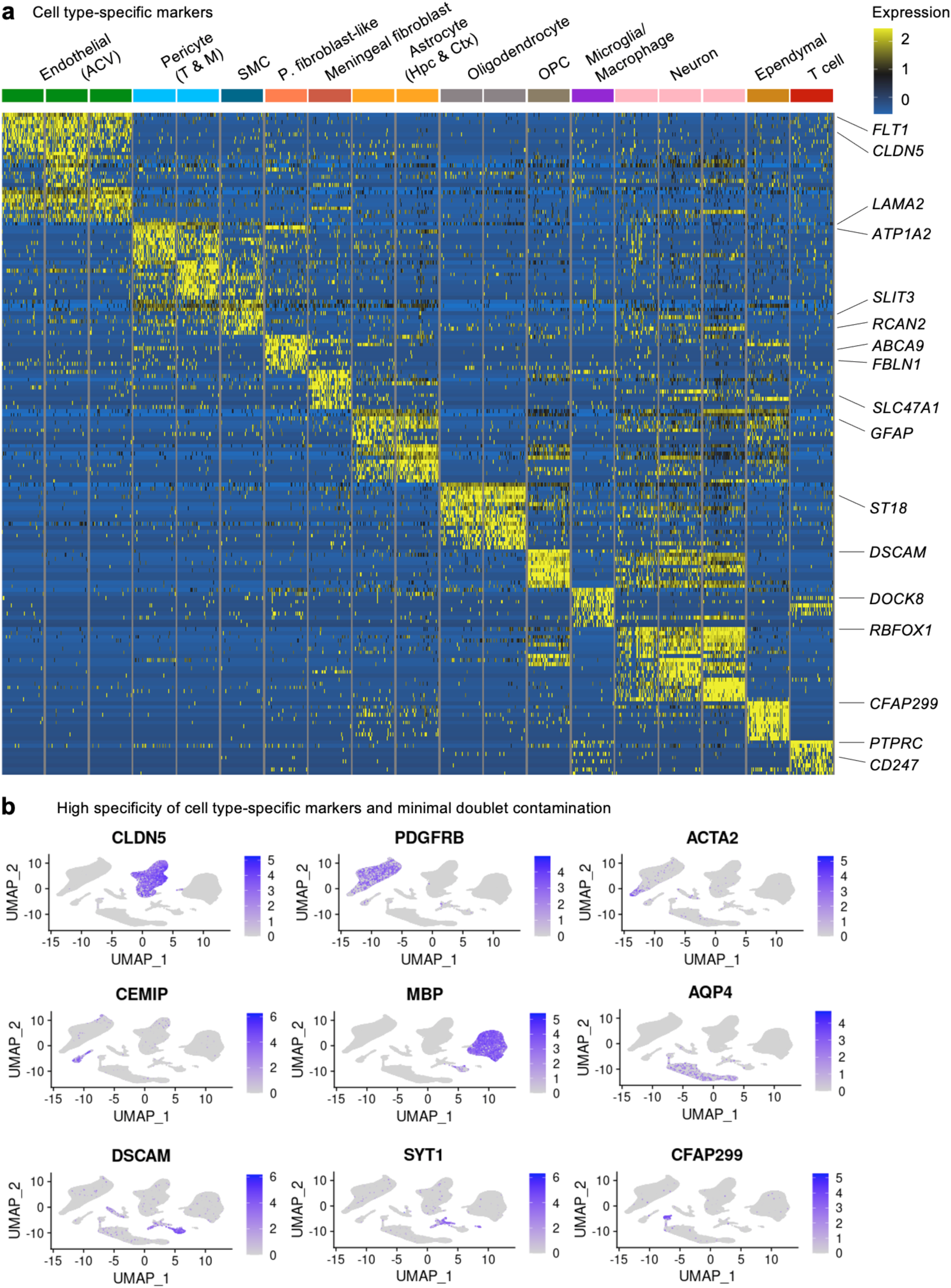
**Cell type-specific gene markers to annotate human cerebrovascular cells.** **a**, Discovery of the top cell type-specific marker genes across the major classes of cells captured. The color bar indicates gene expression from low (blue) to high (yellow). **b**, Validation of cell type annotations and confirmation of minimal doublet contamination using established cell type-specific markers.

**Supplemental Figure 4.**
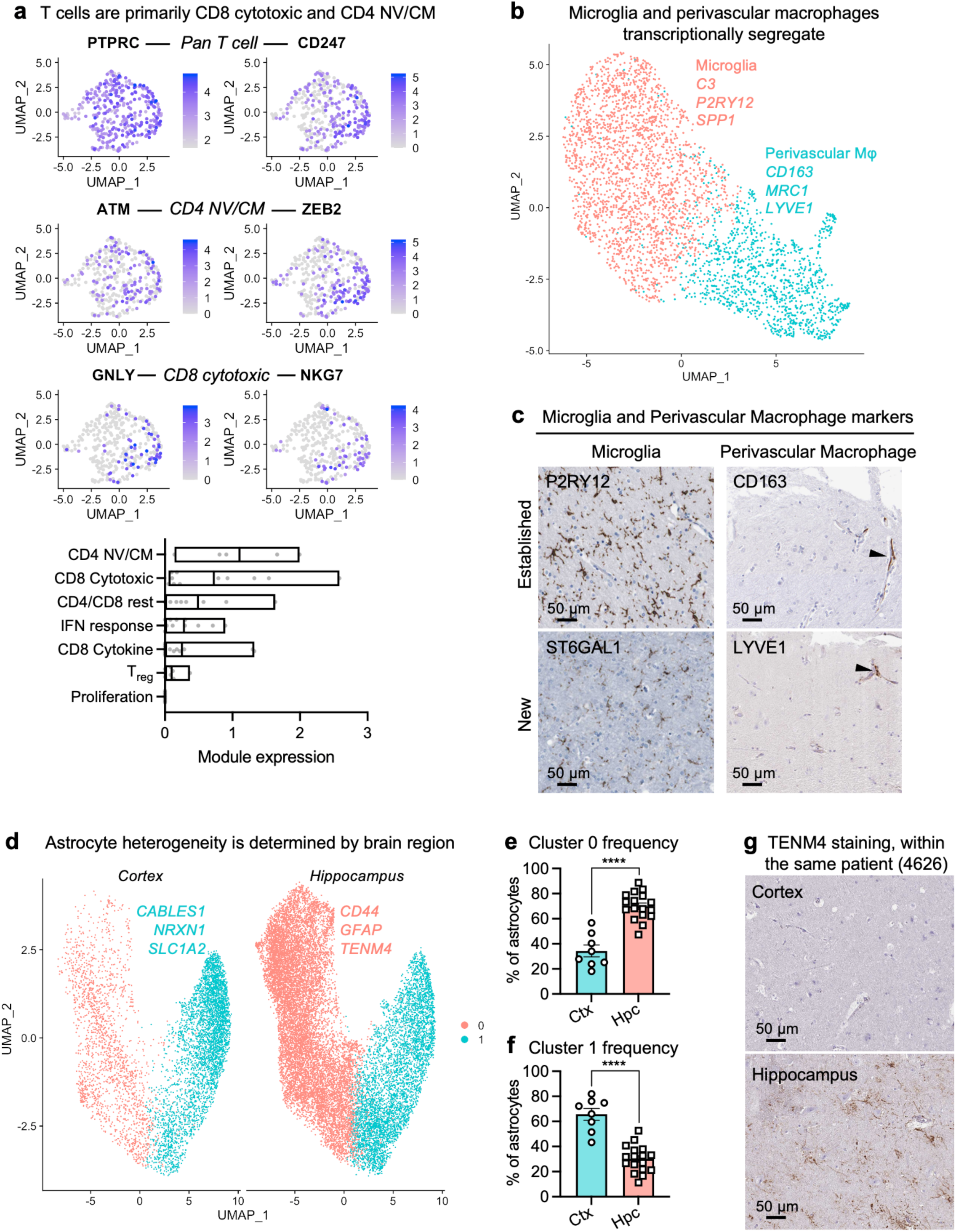
**Perivascular immune cell identity and astrocyte heterogeneity.** **a**, Expression of top gene markers for various T cell subtypes (top), and quantification of their expression as a module (bottom)^154^. Brain perivascular T cells exhibit highest expression of markers corresponding to CD8 cytotoxic and CD4 Naive/Central memory (NV/CM) T cells. **b**, UMAP projection of captured myeloid cells, forming two distinct clusters corresponding to parenchymal microglia and perivascular macrophages. Example marker genes listed. Positive perivascular macrophage staining denoted by arrowheads. **c**, Immunohistochemical validation of microglial and perivascular macrophage markers. Scale bar = 50 microns. Images from the Human Protein Atlas (http://www.proteinatlas.org)^75, 140^. **d**, UMAP projection of captured astrocytes, forming two distinct clusters, and split by brain region. Example marker genes listed. **e**-**f**, Quantification of astrocyte cluster 0 (**b**) and 1 (**c**) frequency in the cortex and hippocampus (*n*=8 cortex and *n*=17 hippocampus, Mann-Whitney t-test; mean +/- s.e.m.). **g**, Immunohistochemical validation of the brain region-specific astrocyte marker *TENM4*. Scale bar = 50 microns. Images from the Human Protein Atlas (http://www.proteinatlas.org)^75, 140^.

**Supplemental Figure 5.**
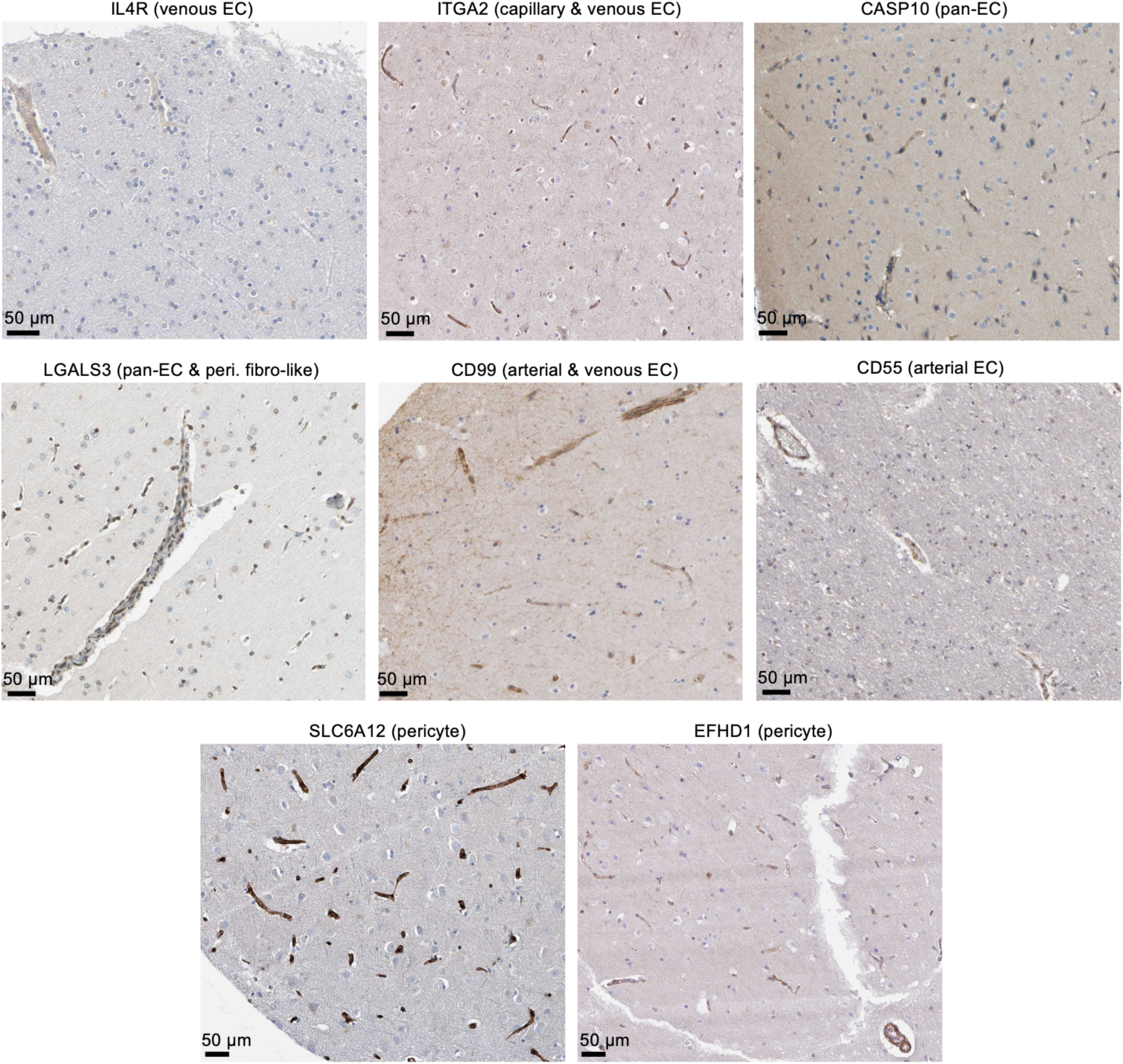
**Examples of predicted human-enriched vascular-expressed genes confirmed *in situ*.** Immunohistochemical confirmation of genes predicted to be enriched or specific to human cerebrovascular cells compared to mouse (isolated mouse nuclei and per Vanlandewijck, et al., 2018)^37^, in terms of overall expression or zonation. In parenthesis is the cell type predicted to be uniquely or exhibiting enriched expressed in human over mouse. Scale bar = 50 microns. Images from the Human Protein Atlas (http://www.proteinatlas.org)^75, 140^.

**Supplemental Figure 6.**
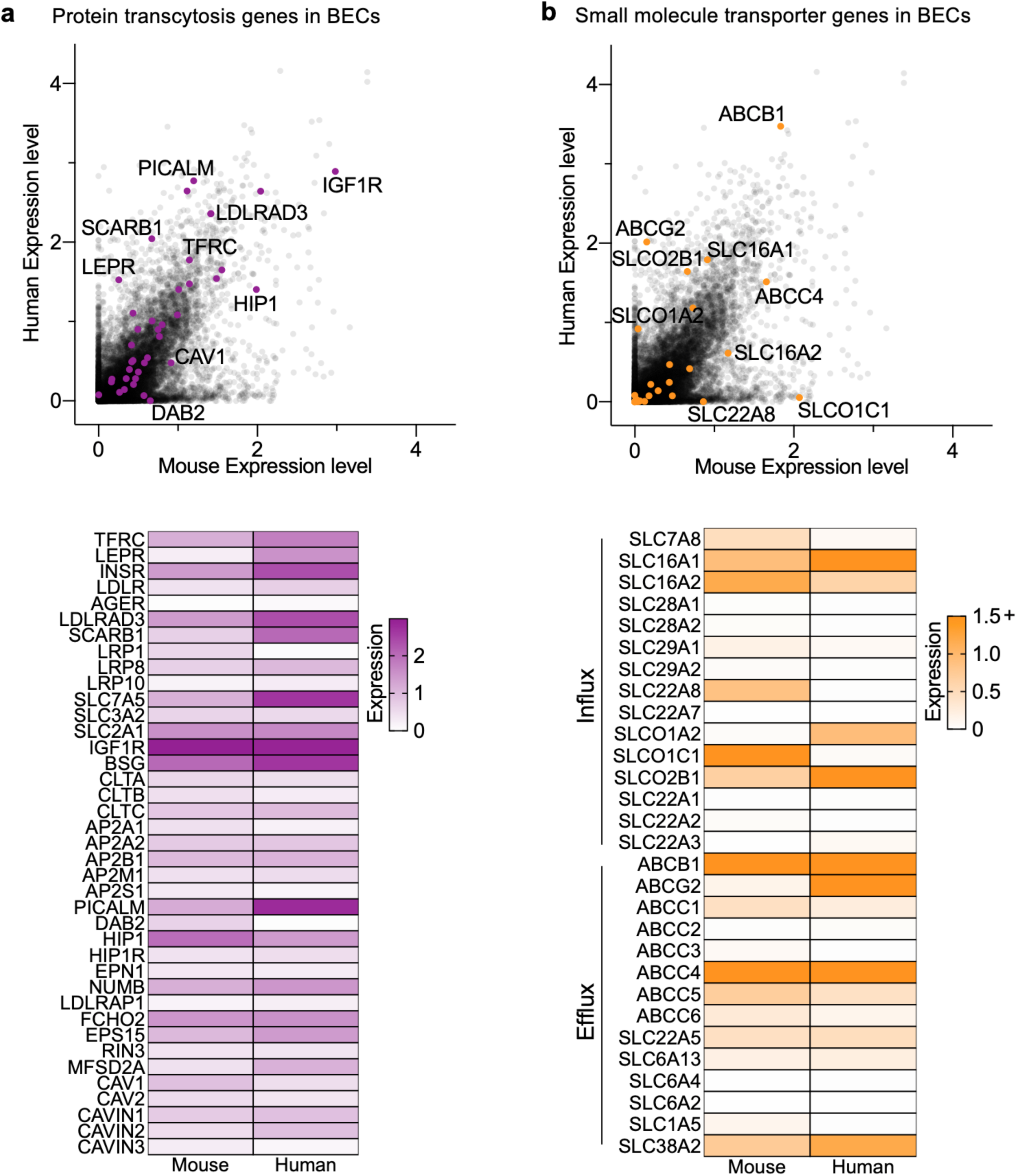
**Comparison of protein transcytosis and small molecule transport genes in mice and human BECs.** Mouse and human brain endothelial cell expression of genes mediating protein transcytosis (**a**) and small molecule influx and efflux (**b**).

**Supplemental Figure 7.**
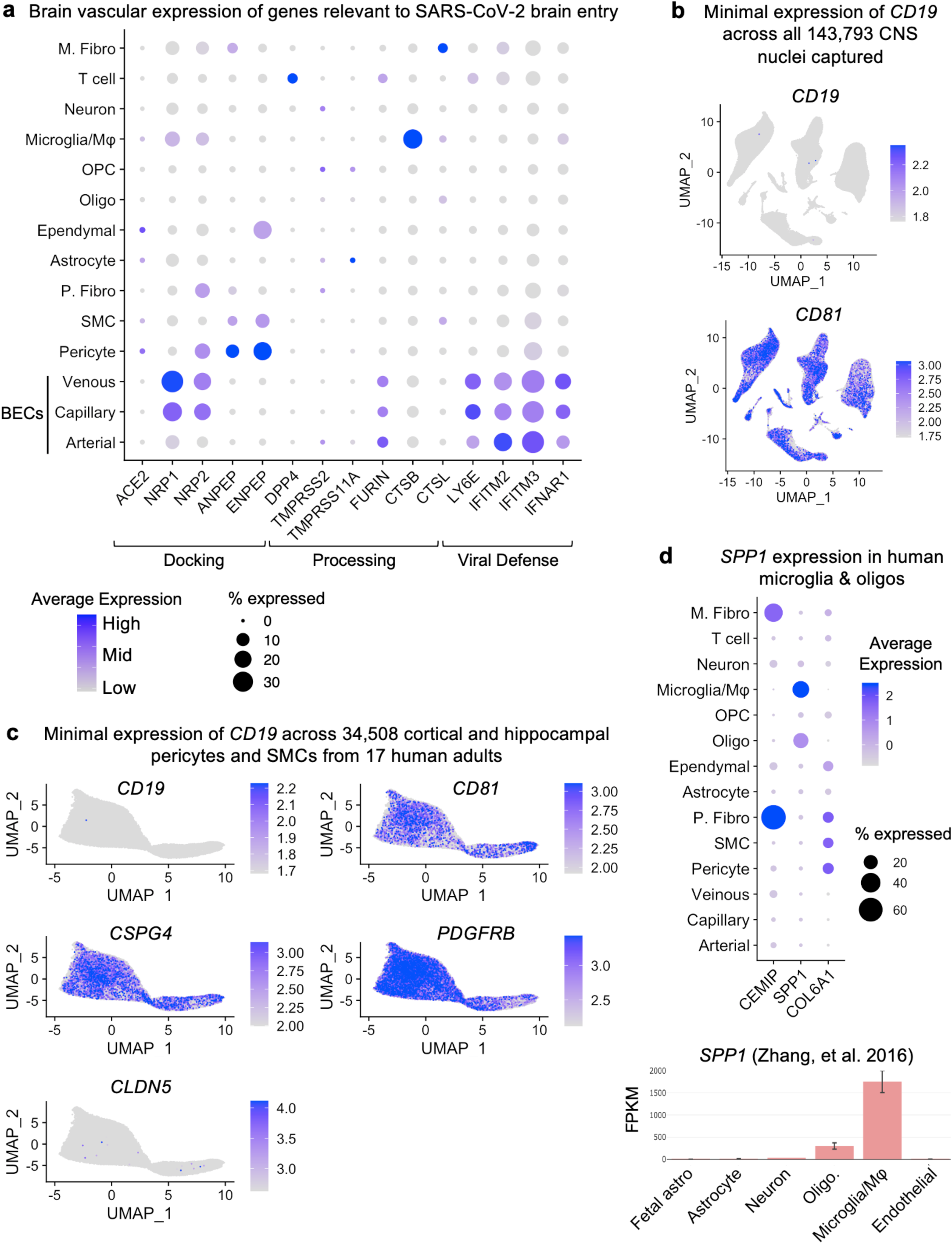
**BBB expression of species-divergent genes relevant to disease.** **a**, Brain vascular expression of genes relevant to SARS-CoV-2 brain entry, as summarized in Iadecola, et al. 2020^66^. **b**-**c**, Expression of the immuno-oncology target *CD19* and its chaperone *CD81* across all 143,793 nuclei captured in this study (**b**) and in human adult brain pericytes and smooth muscle cells (**c**). *Note:* cells with any finite expression are ordered to the front to ensure all expression is visible, but this carries the potential to visually overestimate average expression **d**, Expression of the mouse perivascular fibroblast-like gene *Spp1* is instead specifically expressed in human myeloid cells and oligodendrocytes (*SPP1*, top), with corroborating data from an independent dataset^60^ (bottom).

**Supplemental Figure 8.**
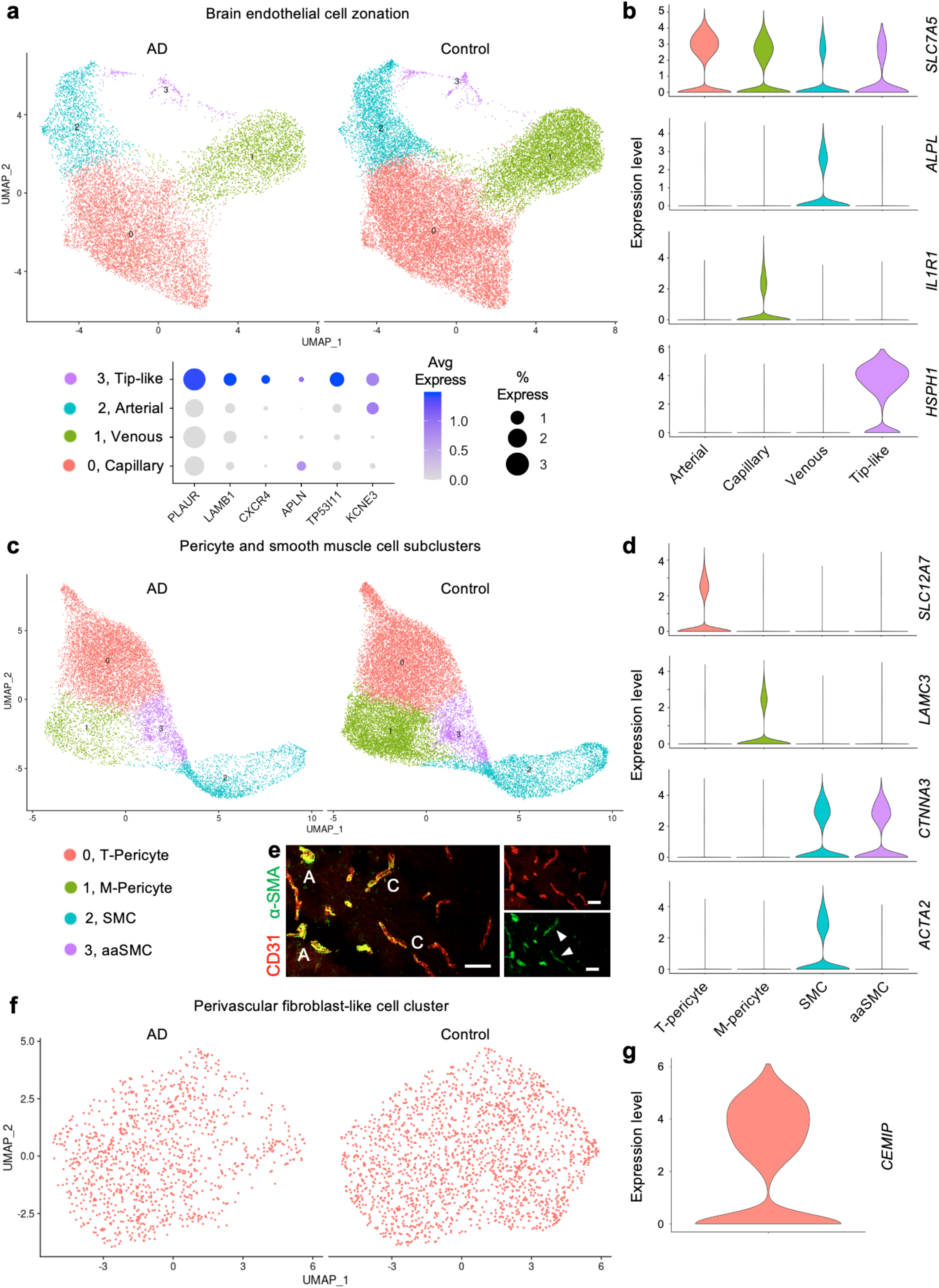
**Brain endothelial and mural cell zonation and subpopulations.** **a**, UMAP projection of captured brain endothelial cells, organizing by arteriovenous zonation. Bottom, tip cell markers expressed in the tip-like/ proteostatic EC cluster. **b**, Validation of brain endothelial cell zonation clusters using established zonation markers^90^. Violin plots are centered around the median, with their shape representing cell distribution. **c**, **d**, As in (**a**-**b**) but for pericytes and smooth muscle cells. Note that the anatomical locations of pericyte 0 and 1 have not yet been determined. **e**, Immunohistochemical validation of *ACTA2* (α-SMA) expression in human smooth muscle cells and less so in capillary pericytes. A denotes arterial and C denotes capillary. Arrowheads specify capillary pericytes expressing *ACTA2*. Scale bar = 50 microns. **f**, **g**, As in (**a**-**b**) but for perivascular fibroblast-like cells, as recently discovered in mice^90^.

**Supplemental Figure 9.**
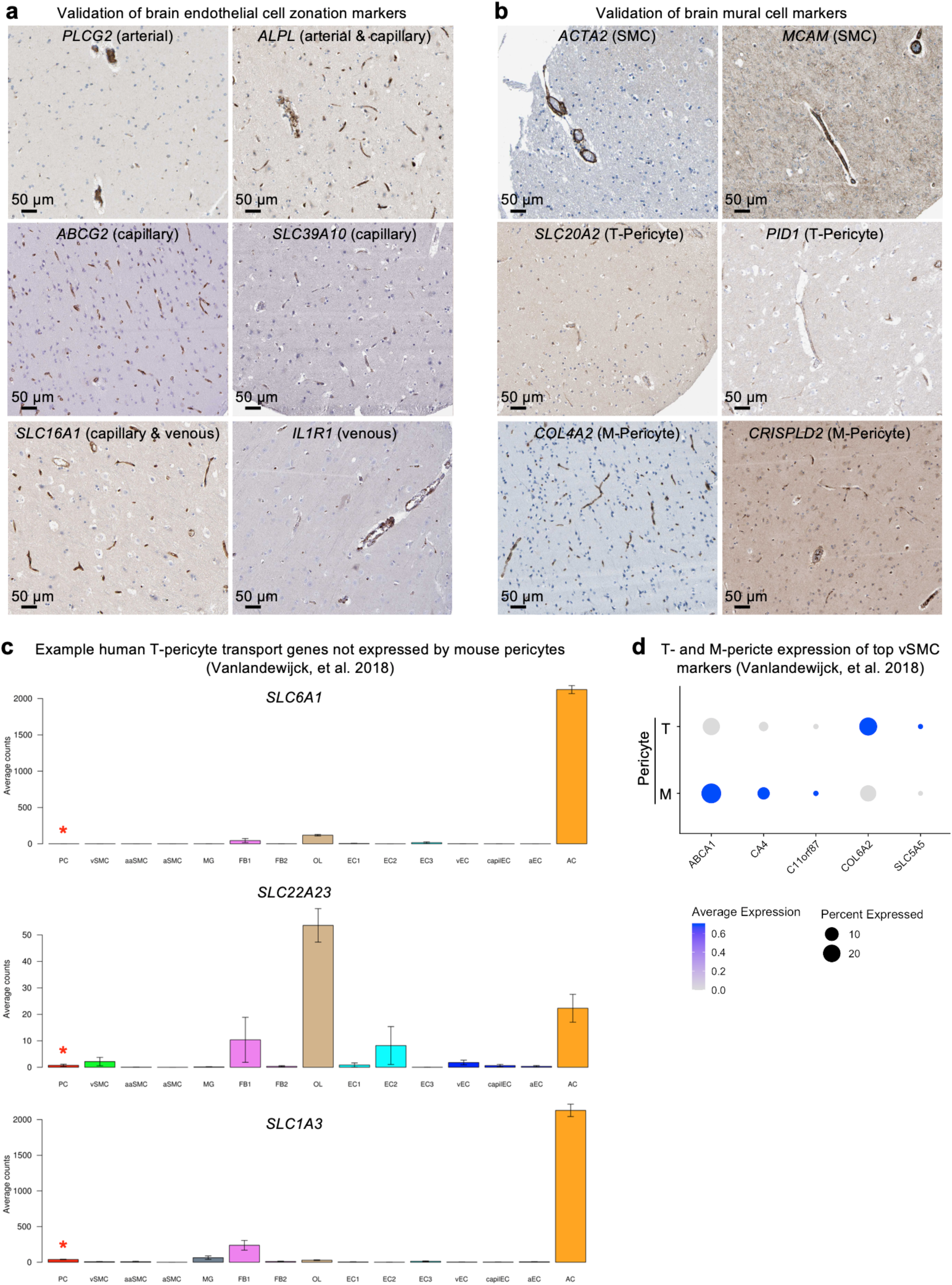
**Validation of brain endothelial and mural cell zonation markers.** **a-b**, Immunohistochemical validation of zonation and cell subtype markers in brain endothelial (**a**) and mural cells (**b**). Scale bar = 50 microns. Images from the Human Protein Atlas (http://www.proteinatlas.org)^75, 140^. **c**, Mouse expression^37^ of example transport genes observed in human T-pericytes, indicating an evolutionary divergence. **d**, Expression of top vSMC markers^37^ across both T- and M-pericytes.

**Supplemental Figure 10.**
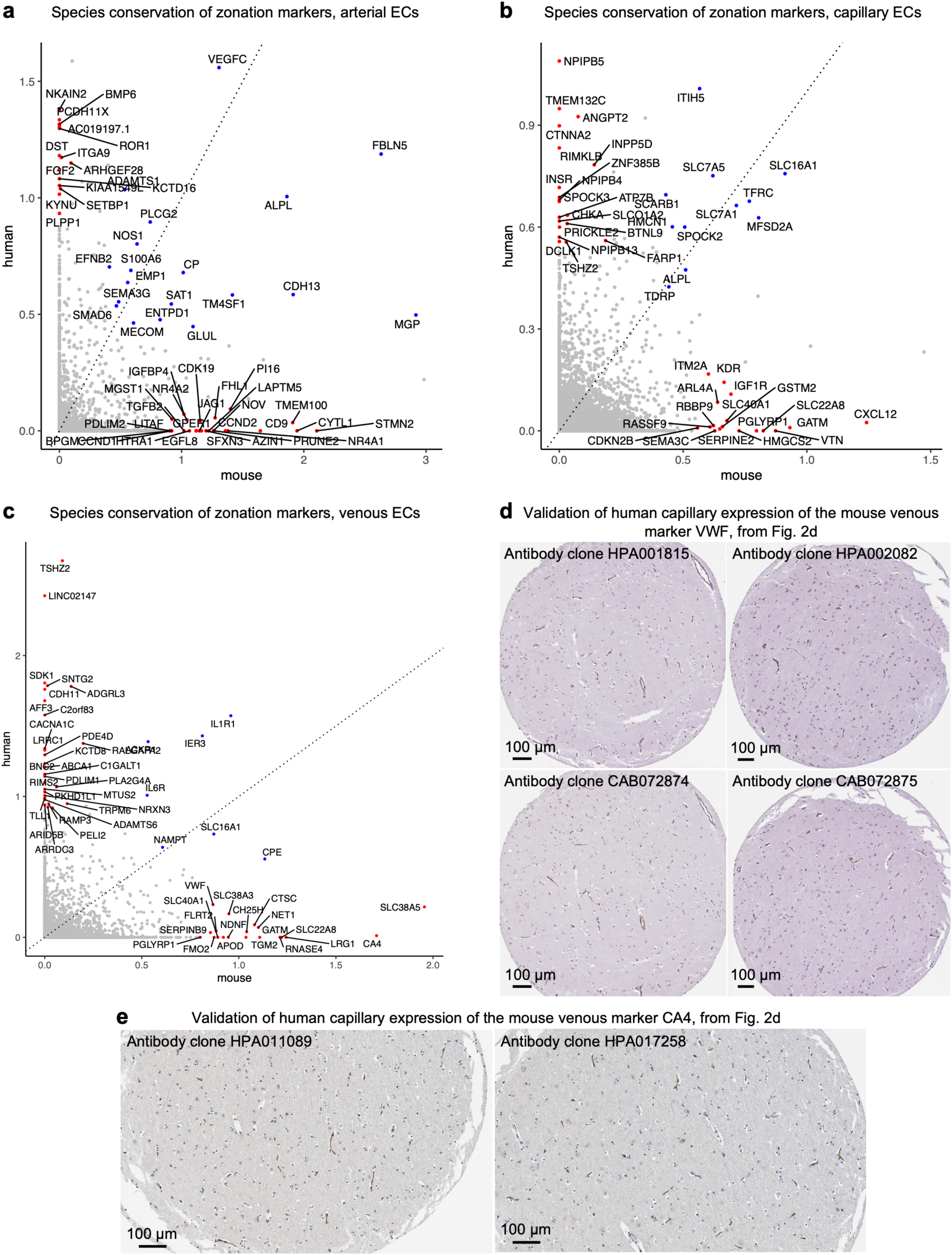
**Zonation markers in mice and humans.** **a-c**, Comparison of the zonal specificity of genes in arterial (**a**), capillary (**b**), and venous (**c**) cells. Axis plot a specificity score, as defined in the Methods. For example, specificity score for capillaries = avg(logFC(cap/ven), logFC(cap/art)) **d-e**, Immunohistochemical validation of capillary expression in human brains of the mouse venous-specific marker VWF (**d**) and CA4 (**e**), with similar patterns observed across multiple primary antibody clones. Scale bar = 100 microns. Images from the Human Protein Atlas (http://www.proteinatlas.org)^75, 140^.

**Supplemental Figure 11.**
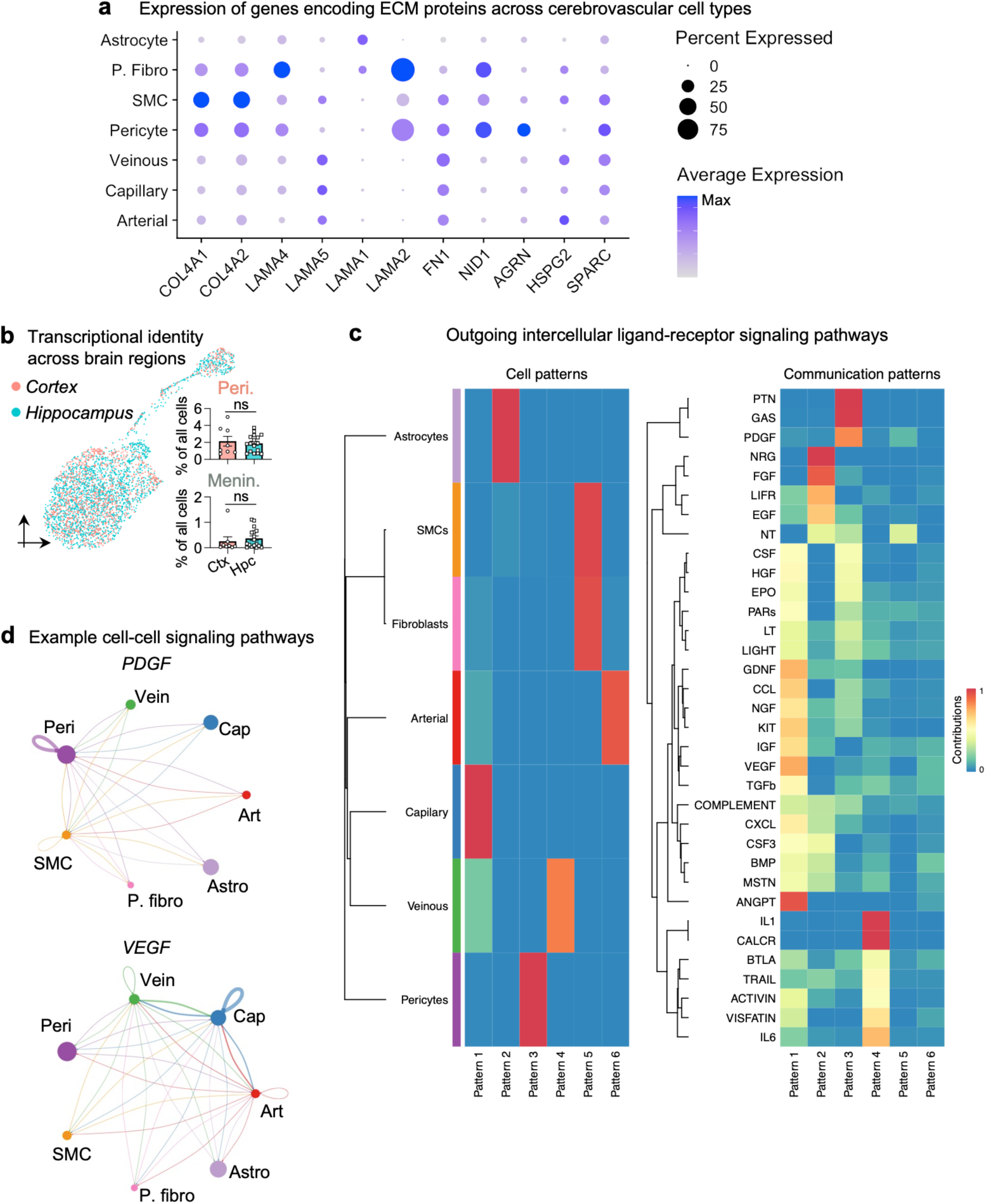
**Extracellular and intercellular interactions in the human BBB.** **a**, Brain vascular expression of genes encoding extracellular matrix proteins, as summarized in Baeten & Akassoglou, 2012^155^. **b**, UMAP of fibroblasts colored by brain region of origin (left). Quantification of perivascular and meningeal fibroblast frequencies from each brain region (right, Mann-Whitney test; mean +/- s.e.m.). **c**, Patterns of outgoing signals from cerebrovascular cells, as identified by ligand-receptor mapping (permutation test, CellChat^93^). **d**, Circle plot showing the number of statistically significant intercellular signaling interactions among cerebrovascular cells for the PDGF and VEGF family of molecules (permutation test, CellChat^93^).

**Supplemental Figure 12.**
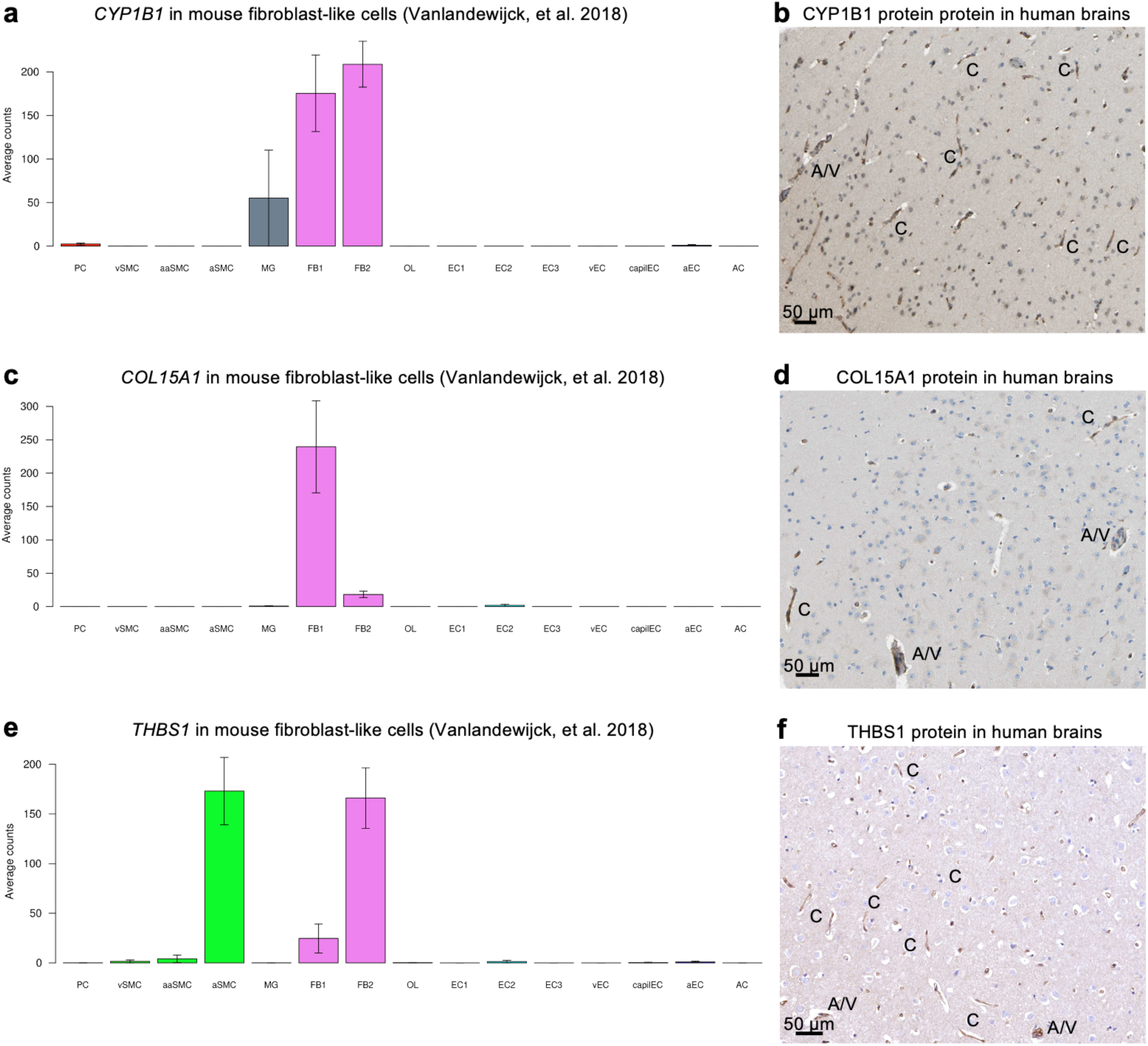
**Proteins specific to perivascular fibroblast-like cells co-localize with human brain capillaries.** **a**, *CYP1B1* expression specifically in mouse^37^ perivascular fibroblast-like cells. **b**, Immunohistochemical localization of fibroblast-derived CYP1B1 with larger diameter arterial and venous (A/V) vasculature as well as capillaries (C) in the human brain. Scale bar = 50 microns. Images from the Human Protein Atlas (http://www.proteinatlas.org)^75, 140^. **c-f**, As in (**a-b**) but for other fibroblast-enriched genes, COL15A1 (**c**-**d**) and THBS1 (**e**-**f**).

**Supplemental Figure 13.**
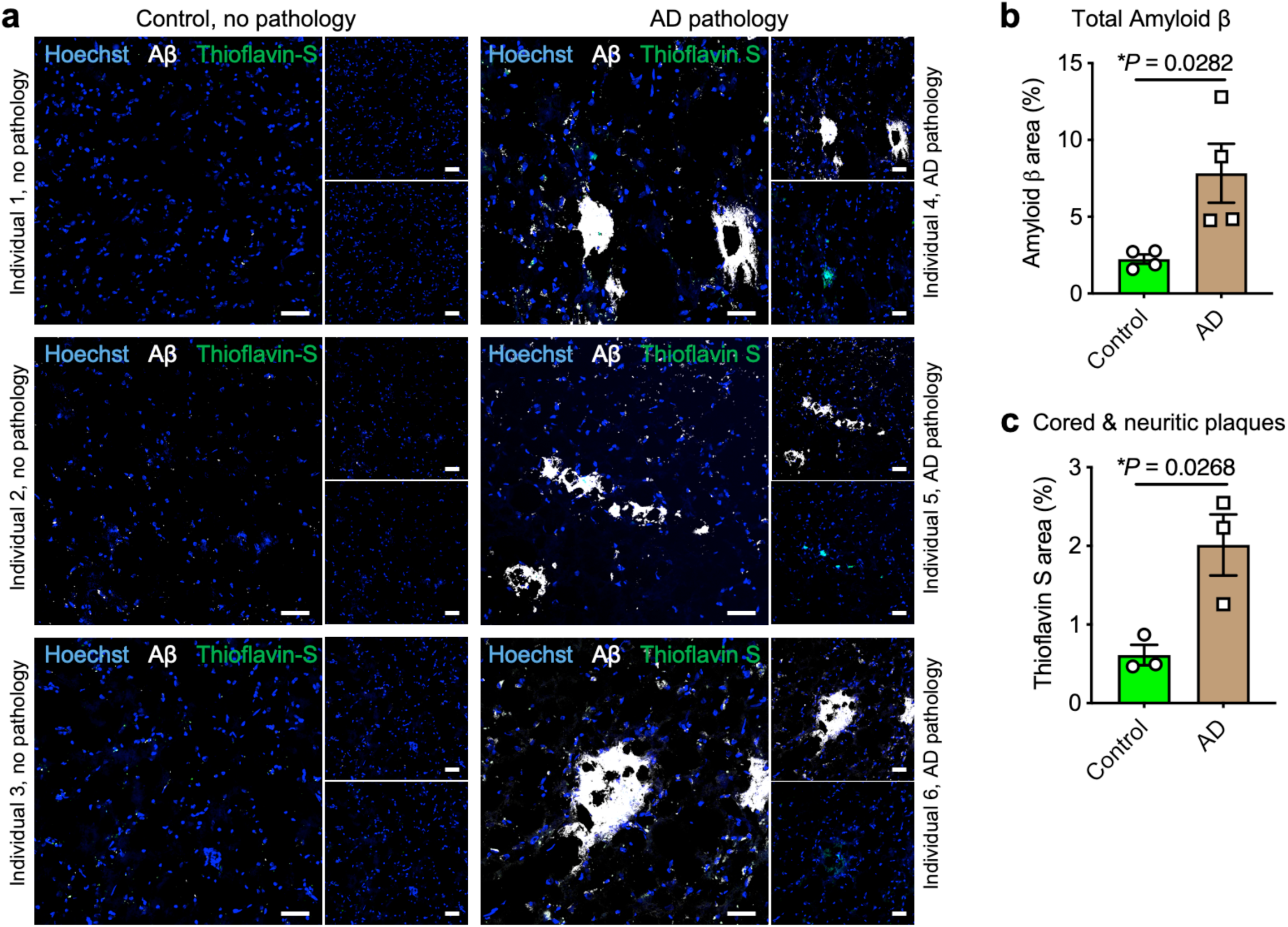
**Confirmation of AD pathology**. **a**, Immunohistochemistry with anti-β-amyloid antibody (D54D2, white), Thioflavin S (green), and Hoechst (blue) in the hippocampus of control and AD patients. Scale bar = 40 microns. **b**, Quantification of β-amyloid immunostaining in (**a**) for overall β-amyloid (*n*=4 controls and AD, two-sided t-test; mean +/- s.e.m.). **c**, As in (**b**) but for cored and neuritic β-amyloid plaques (*n*=3 controls and AD, two-sided t-test; mean +/- s.e.m.).

**Supplemental Figure 14.**
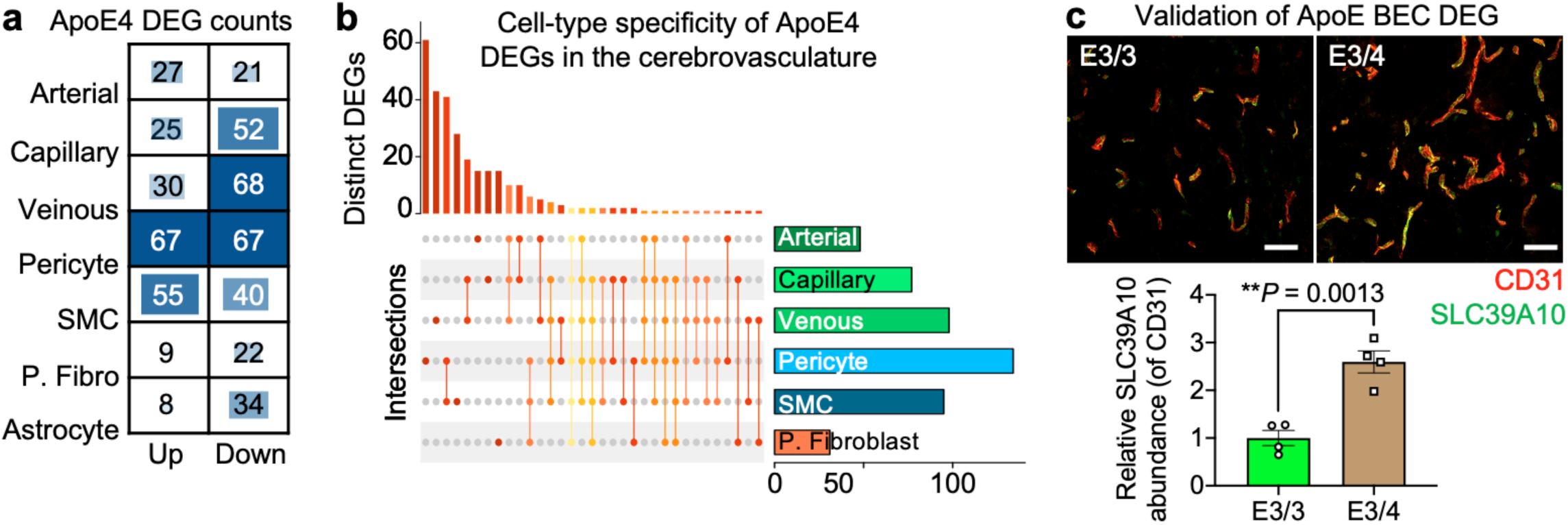
**Vascular cell-type specific perturbations in ApoE4 carriers.** **a**, Differentially expressed gene (DEG) counts for each cell type in ApoE4 carriers (*n* = 5 ApoE3/3, *n* = 11 ApoE3/4 or ApoE4/4): arterial (Art), capillary (Cap), venous (Vein), pericyte (Peri), perivascular fibro blast-like cell (P. fibro), and smooth muscle cell (SMC). The intensity of the blue color and the size of the squares are proportional to entry values. **b**, Matrix layout for intersections of ApoE4 DEGs shared across and specific to each cell type. Circles in the matrix indicate sets that are part of the intersection, showing that most DEGs are cell type-specific. **c**, Immunohistochemical validation of the predicted upregulated anti-inflammatory DEG *SLC39A10* in venous BECs of ApoE4 carriers. Scale bar = 50 microns (*n* = 4 ApoE3/3 and ApoE4 carriers, nested two-sided t-test; mean +/- s.e.m.).

**Supplemental Figure 15.**
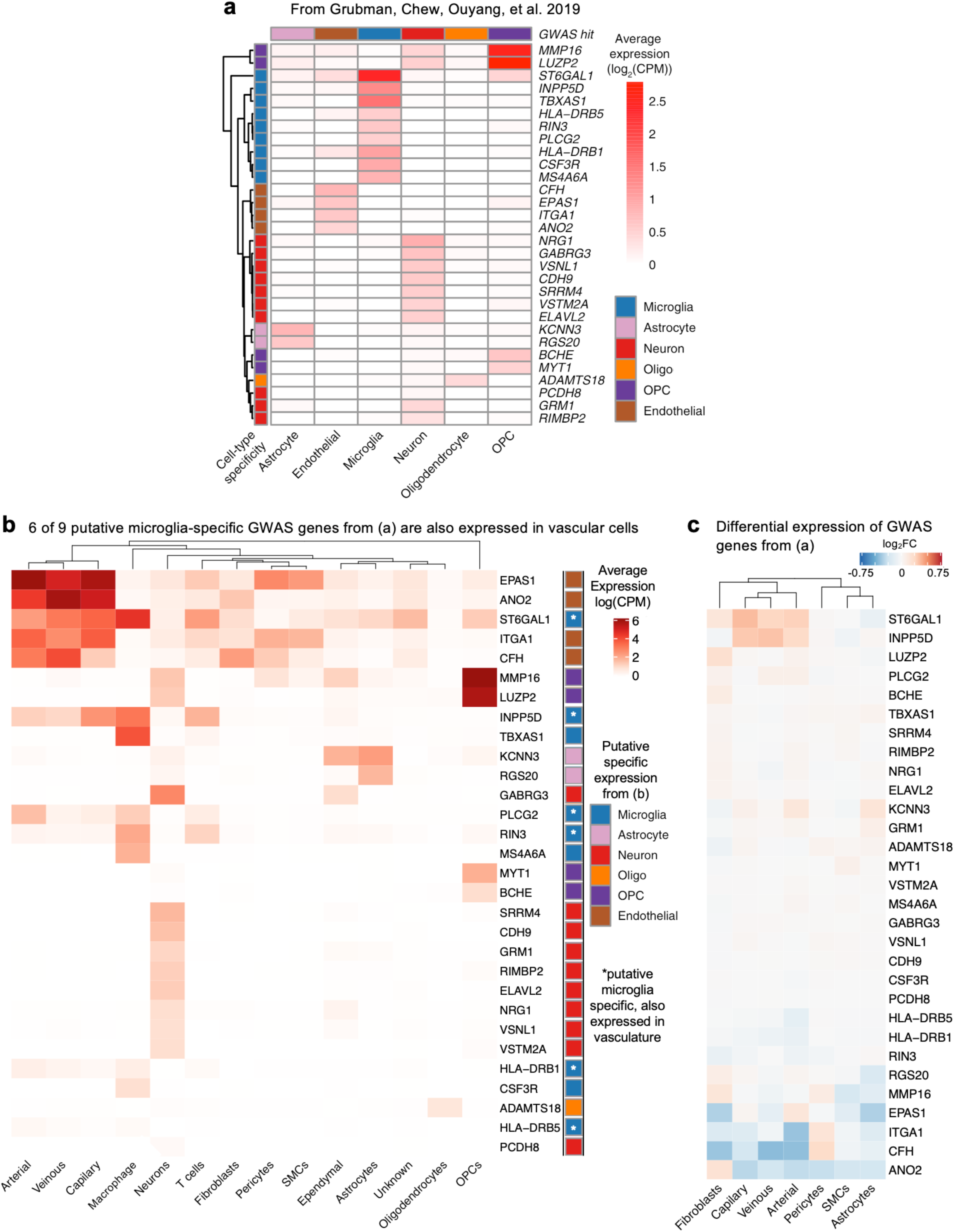
**Re-evaluation of putative cell type-specifically expressed AD GWAS genes.** **a**, GWAS genes found to be expressed cell-type specifically among cells captured using the conventional nuclei isolation process, from Grubman, Chew, Ouyang, et al. 2019^29^. **b**, As in (**a**) but now evaluated for cells captured with the new nuclei isolation process. Several GWAS genes believed specifically expressed in microglia are also expressed in vascular cells (asterisks). **c**, As in (**a**-**b**), but plotting differential expression of these GWAS risk genes amongst vascular cells in AD.

**Supplemental Figure 16.**
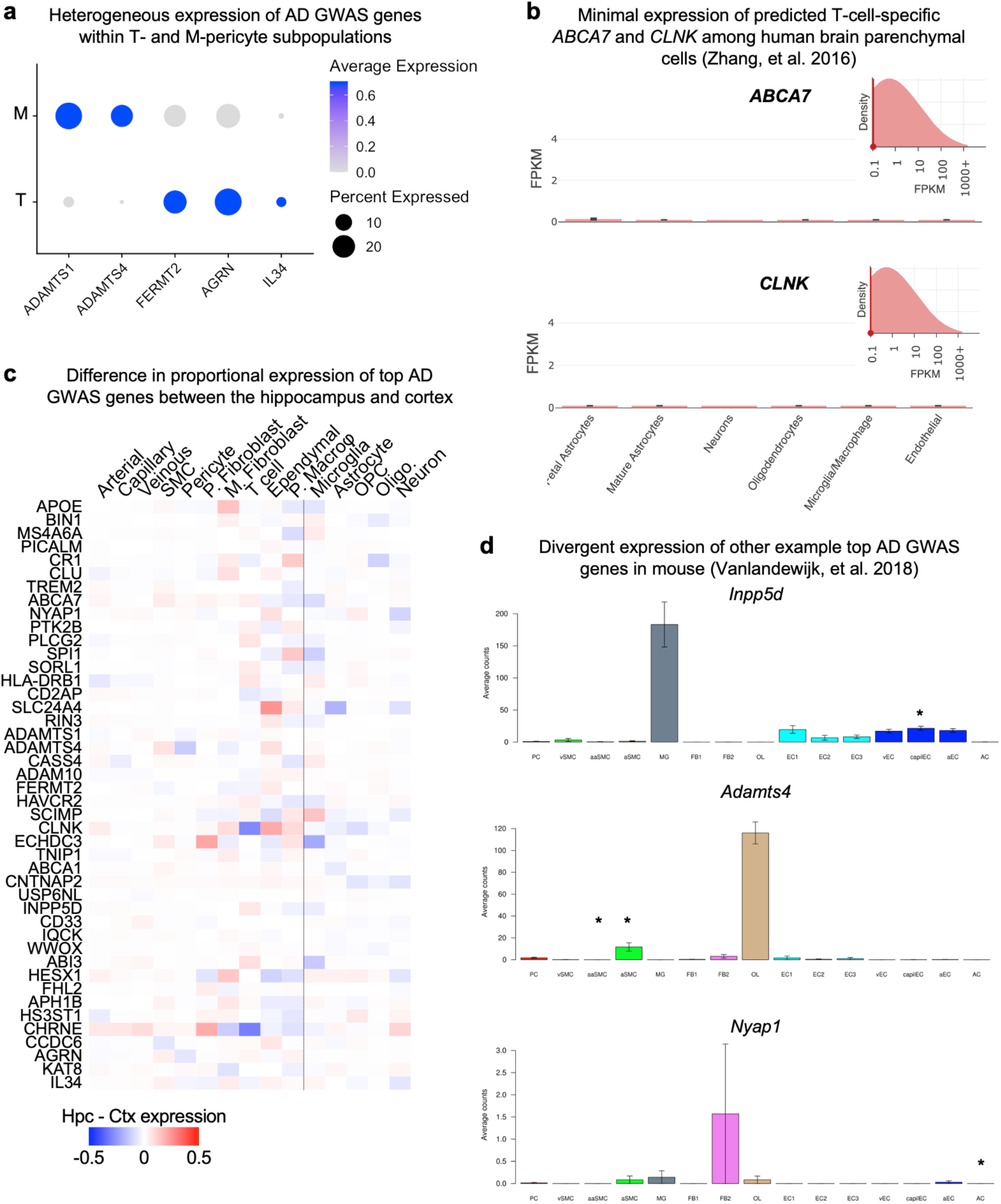
**Brain region- and species-specific expression of top AD GWAS genes.** **a**, Heterogeneous expression of AD GWAS genes across T- and M-pericyte subtypes. **b**, RNA-seq data of the predicted perivascular T cell-specific AD GWAS genes *EPHA1* and *ABCA7* in an independent dataset^60^, corroborating minimal expression across resident/ parenchymal brain cells. **c**, Heatmap comparing expression patterns of top AD GWAS genes in the hippocampus and superior frontal cortex: e.g., several microglia-expressed GWAS genes like *APOE*, *MS4A4A*, and *TREM2* are more highly expressed in hippocampal compared to cortical microglia/ macrophages. **d**, Expression of top AD GWAS genes in mouse^37^. Asterisks indicate vascular cell types where gene is also strongly expressed in humans (Fig. 5a-b).

**Supplemental Figure 17.**
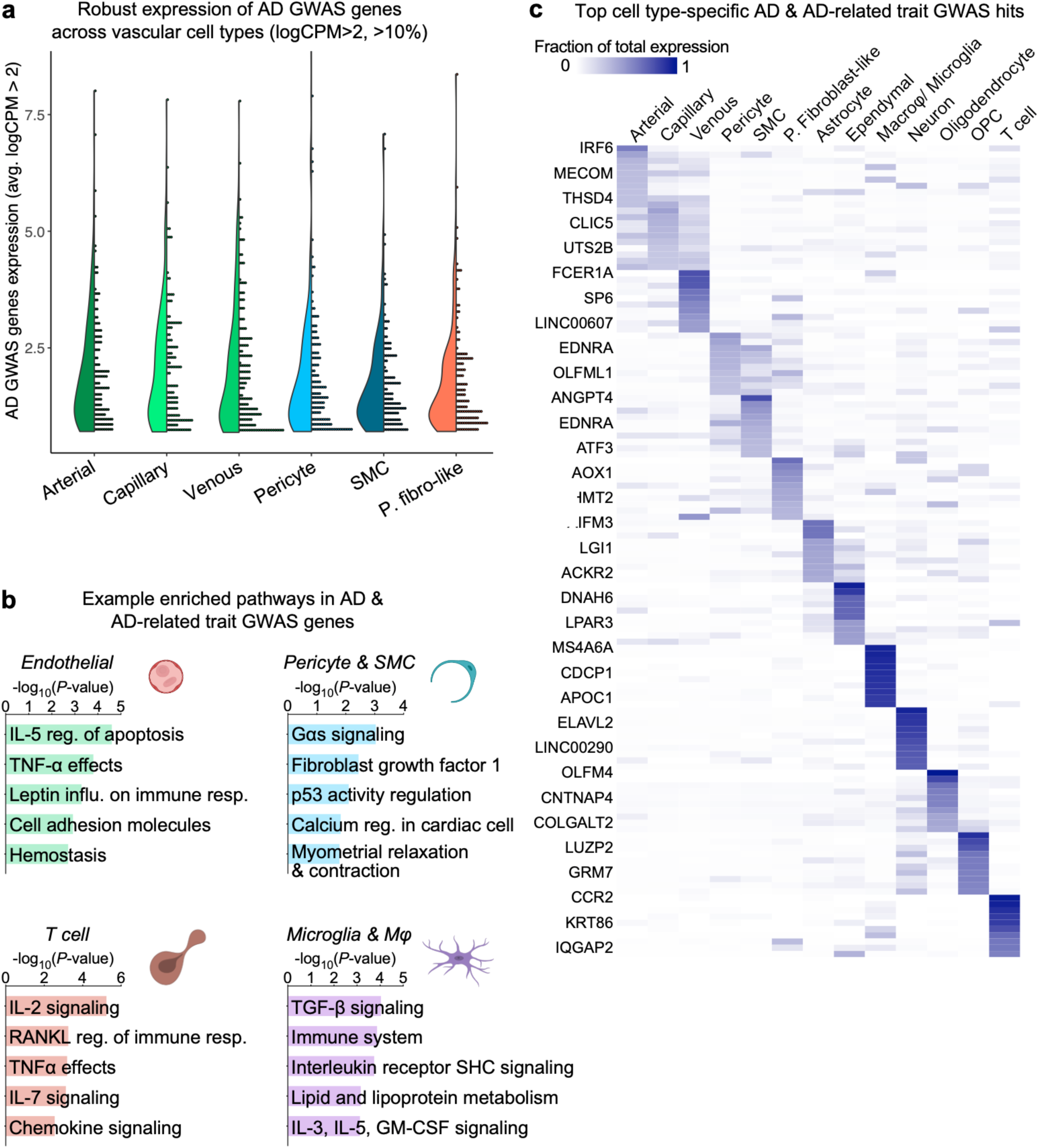
**Brain vascular and perivascular expression of major AD GWAS genes.** **a**, Expression of Alzheimer’s disease (AD) and AD-related GWAS risk genes (from Grubman, Chew, Ouyang, et al. 2019)^29^ across human vascular cells. **b**, Enriched biological pathways amongst AD and AD-related trait GWAS genes expressed in each cell type. **c**, For each cell type, the top 10 most specifically expressed AD and AD-related trait GWAS genes.

## Notes

https://twc-stanford.shinyapps.io/human_bbb

### Competing Interest Statement

The authors have declared no competing interest.

## References

1. Cipolla, M. J. The Cerebral Circulation, Second Edition. Colloq. Ser. Integr. Syst. Physiol. From Mol. to Funct. (2016). doi:10.4199/c00141ed2v01y201607isp066

2. Feigin, V. L. et al. Global and regional burden of stroke during 1990-2010: Findings from the Global Burden of Disease Study 2010. Lancet (2014). doi:10.1016/S0140-6736(13)61953-4

3. Tong, X. et al. The burden of cerebrovascular disease in the United States. Prev. Chronic Dis. (2019). doi:10.5888/pcd16.180411

4. Gorelick, P. B. The global burden of stroke: persistent and disabling. The Lancet Neurology (2019). doi:10.1016/S1474-4422(19)30030-4

5. Zhao, Z., Nelson, A. R., Betsholtz, C. & Zlokovic, B. V. Establishment and Dysfunction of the Blood-Brain Barrier. Cell 163, 1064–1078 (2015).

6. Sweeney, M. D., Zhao, Z., Montagne, A., Nelson, A. R. & Zlokovic, B. V. Blood-brain barrier: From physiology to disease and back. Physiological Reviews (2019). doi:10.1152/physrev.00050.2017

7. Iadecola, C. The Pathobiology of Vascular Dementia. Neuron (2013). doi:10.1016/j.neuron.2013.10.008

8. Zlokovic, B. V. The blood-brain barrier in health and chronic neurodegenerative disorders. Neuron (2008). doi:10.1016/j.neuron.2008.01.003

9. Montagne, A. et al. Blood-Brain barrier breakdown in the aging human hippocampus. Neuron (2015). doi:10.1016/j.neuron.2014.12.032

10. Montagne, A., Zhao, Z. & Zlokovic, B. V. Alzheimer’s disease: A matter of blood–brain barrier dysfunction? J. Exp. Med. (2017). doi:10.1084/jem.20171406

11. Banks, W. A., Reed, M. J., Logsdon, A. F., Rhea, E. M. & Erickson, M. A. Healthy aging and the blood–brain barrier. Nat. Aging 1, (2021).

12. Daneman, R. The blood-brain barrier in health and disease. Ann. Neurol. (2012). doi:10.1002/ana.23648

13. Chow, B. W. & Gu, C. The Molecular Constituents of the Blood-Brain Barrier. Trends Neurosci. 38, 598–608 (2015).

14. Profaci, C. P., Munji, R. N., Pulido, R. S. & Daneman, R. The blood-brain barrier in health and disease: Important unanswered questions. J. Exp. Med. (2020). doi:10.1084/jem.20190062

15. Yang, A. C. et al. Physiological blood–brain transport is impaired with age by a shift in transcytosis. Nature (2020). doi:10.1038/s41586-020-2453-z

16. Obermeier, B., Daneman, R. & Ransohoff, R. M. Development, maintenance and disruption of the blood-brain barrier. Nature Medicine (2013). doi:10.1038/nm.3407

17. Abbott, N. J., Rönnbäck, L. & Hansson, E. Astrocyte-endothelial interactions at the blood-brain barrier. Nature Reviews Neuroscience (2006). doi:10.1038/nrn1824

18. Pardridge, W. M. Drug transport across the blood-brain barrier. Journal of Cerebral Blood Flow and Metabolism (2012). doi:10.1038/jcbfm.2012.126

19. de Lange, E. C. M. & Hammarlund-Udenaes, M. Translational aspects of blood-brain barrier transport and central nervous system effects of drugs: from discovery to patients. Clinical pharmacology and therapeutics (2015). doi:10.1002/cpt.76

20. Zuchero, Y. J. Y. et al. Discovery of Novel Blood-Brain Barrier Targets to Enhance Brain Uptake of Therapeutic Antibodies. Neuron (2016). doi:10.1016/j.neuron.2015.11.024

21. Yu, Y. J. & Watts, R. J. Developing Therapeutic Antibodies for Neurodegenerative Disease. Neurotherapeutics (2013). doi:10.1007/s13311-013-0187-4

22. Kariolis, M. S. et al. Brain delivery of therapeutic proteins using an Fc fragment blood-brain barrier transport vehicle in mice and monkeys. Sci. Transl. Med. 12, (2020).

23. Daneman, R., Zhou, L., Kebede, A. A. & Barres, B. A. Pericytes are required for bloodĝ€”brain barrier integrity during embryogenesis. Nature 468, 562–566 (2010).

24. Armulik, A. et al. Pericytes regulate the blood-brain barrier. Nature 468, 557–561 (2010).

25. Villabona-Rueda, A., Erice, C., Pardo, C. A. & Stins, M. F. The Evolving Concept of the Blood Brain Barrier (BBB): From a Single Static Barrier to a Heterogeneous and Dynamic Relay Center. Frontiers in Cellular Neuroscience (2019). doi:10.3389/fncel.2019.00405

26. Wilhelm, I., Nyúl-Tóth, Á., Suciu, M., Hermenean, A. & Krizbai, I. A. Heterogeneity of the blood-brain barrier. Tissue Barriers (2016). doi:10.1080/21688370.2016.1143544

27. Sweeney, M. D., Kisler, K., Montagne, A., Toga, A. W. & Zlokovic, B. V. The role of brain vasculature in neurodegenerative disorders. Nature Neuroscience (2018). doi:10.1038/s41593-018-0234-x

28. Mathys, H. et al. Single-cell transcriptomic analysis of Alzheimer’s disease. Nature (2019). doi:10.1038/s41586-019-1195-2

29. Grubman, A. et al. A single-cell atlas of entorhinal cortex from individuals with Alzheimer’s disease reveals cell-type-specific gene expression regulation. Nat. Neurosci. (2019). doi:10.1038/s41593-019-0539-4

30. Zhou, Y. et al. Human and mouse single-nucleus transcriptomics reveal TREM2-dependent and TREM2-independent cellular responses in Alzheimer’s disease. Nat. Med. (2020). doi:10.1038/s41591-019-0695-9

31. Jäkel, S. et al. Altered human oligodendrocyte heterogeneity in multiple sclerosis. Nature (2019). doi:10.1038/s41586-019-0903-2

32. Velmeshev, D. et al. Single-cell genomics identifies cell type–specific molecular changes in autism. Science (80-.). (2019). doi:10.1126/science.aav8130

33. Al-Dalahmah, O. et al. Single-nucleus RNA-seq identifies Huntington disease astrocyte states. Acta Neuropathol. Commun. (2020). doi:10.1186/s40478-020-0880-6

34. Keller, D., Erö, C. & Markram, H. Cell densities in the mouse brain: A systematic review. Frontiers in Neuroanatomy (2018). doi:10.3389/fnana.2018.00083

35. Niedowicz, D. M. et al. Obesity and diabetes cause cognitive dysfunction in the absence of accelerated β-amyloid deposition in a novel murine model of mixed or vascular dementia. Acta Neuropathol. Commun. (2014). doi:10.1186/2051-5960-2-64

36. Geisert, E. E. et al. Increased brain size and glial cell number in CD81-null mice. J. Comp. Neurol. (2002). doi:10.1002/cne.10364

37. Vanlandewijck, M. et al. A molecular atlas of cell types and zonation in the brain vasculature. Nature 554, 475–480 (2018).

38. Sabbagh, M. F. et al. Transcriptional and epigenomic landscapes of CNS and non-CNS vascular endothelial cells. Elife (2018). doi:10.7554/elife.36187

39. Kalucka, J. et al. Single-Cell Transcriptome Atlas of Murine Endothelial Cells. Cell (2020). doi:10.1016/j.cell.2020.01.015

40. Schaum, N. et al. Single-cell transcriptomics of 20 mouse organs creates a Tabula Muris. Nature (2018). doi:10.1038/s41586-018-0590-4

41. Chen, M. B. et al. Brain endothelial cells are exquisite sensors of age-related circulatory cues. bioRxiv (2019). doi:10.1101/617258

42. Saunders, A. et al. Molecular Diversity and Specializations among the Cells of the Adult Mouse Brain. Cell (2018). doi:10.1016/j.cell.2018.07.028

43. Nei, M., Xu, P. & Glazko, G. Estimation of divergence times from multiprotein sequences for a few mammalian species and several distantly related organisms. Proc. Natl. Acad. Sci. U. S. A. (2001). doi:10.1073/pnas.051611498

44. Geirsdottir, L. et al. Cross-Species Single-Cell Analysis Reveals Divergence of the Primate Microglia Program. Cell (2019). doi:10.1016/j.cell.2019.11.010

45. Huang, Q. et al. Delivering genes across the blood-brain barrier: LY6A, a novel cellular receptor for AAV-PHP.B capsids. PLoS One (2019). doi:10.1371/journal.pone.0225206

46. Batista, A. R. et al. Ly6a Differential Expression in Blood-Brain Barrier Is Responsible for Strain Specific Central Nervous System Transduction Profile of AAV-PHP.B. Hum. Gene Ther. (2020). doi:10.1089/hum.2019.186

47. Hordeaux, J. et al. The GPI-Linked Protein LY6A Drives AAV-PHP.B Transport across the Blood-Brain Barrier. Mol. Ther. (2019). doi:10.1016/j.ymthe.2019.02.013

48. Corces, M. R. et al. Single-cell epigenomic analyses implicate candidate causal variants at inherited risk loci for Alzheimer’s and Parkinson’s diseases. Nat. Genet. (2020). doi:10.1038/s41588-020-00721-x

49. Habib, N. et al. Massively parallel single-nucleus RNA-seq with DroNc-seq. Nat. Methods (2017). doi:10.1038/nmeth.4407

50. Litviňuková, M. et al. Cells of the adult human heart. Nature (2020). doi:10.1038/s41586-020-2797-4

51. Leng, K. et al. Molecular characterization of selectively vulnerable neurons in Alzheimer’s disease. Nat. Neurosci. (2021). doi:10.1038/s41593-020-00764-7

52. Corces, M. R. et al. An improved ATAC-seq protocol reduces background and enables interrogation of frozen tissues. Nat. Methods (2017). doi:10.1038/nmeth.4396

53. Triguero, D., Buciak, J. & Pardridge, W. M. Capillary Depletion Method for Quantification of Blood–Brain Barrier Transport of Circulating Peptides and Plasma Proteins. J. Neurochem. (1990). doi:10.1111/j.1471-4159.1990.tb04886.x

54. Lee, Y. K., Uchida, H., Smith, H., Ito, A. & Sanchez, T. The isolation and molecular characterization of cerebral microvessels. Nat. Protoc. (2019). doi:10.1038/s41596-019-0212-0

55. Kruisbeek, A. M. Isolation of Mouse Mononuclear Cells. Curr. Protoc. Immunol. (2000). doi:10.1002/0471142735.im0301s39

56. Ong, W. Y. & Levine, J. M. A light and electron microscopic study of NG2 chondroitin sulfate proteoglycan-positive oligodendrocyte precursor cells in the normal and kainate-lesioned rat hippocampus. Neuroscience (1999). doi:10.1016/S0306-4522(98)00751-9

57. Dulken, B. W. et al. Single-cell analysis reveals T cell infiltration in old neurogenic niches. Nature (2019). doi:10.1038/s41586-019-1362-5

58. Yousef, H. et al. Aged blood impairs hippocampal neural precursor activity and activates microglia via brain endothelial cell VCAM1. Nat. Med. (2019). doi:10.1038/s41591-019-0440-4

59. Smith, L. K. et al. Β2-Microglobulin Is a Systemic Pro-Aging Factor That Impairs Cognitive Function and Neurogenesis. Nat. Med. 21, 932–937 (2015).

60. Zhang, Y. et al. Purification and Characterization of Progenitor and Mature Human Astrocytes Reveals Transcriptional and Functional Differences with Mouse. Neuron (2016). doi:10.1016/j.neuron.2015.11.013

61. Maier, M. et al. Complement C3 deficiency leads to accelerated amyloid β plaque deposition and neurodegeneration and modulation of the microglia/macrophage phenotype in amyloid precursor protein transgenic mice. J. Neurosci. (2008). doi:10.1523/JNEUROSCI.0829-08.2008

62. Hughes, S. R. et al. α2-macroglobulin associates with β-amyloid peptide and prevents fibril formation. Proc. Natl. Acad. Sci. U. S. A. (1998). doi:10.1073/pnas.95.6.3275

63. Beck, T. N., Nicolas, E., Kopp, M. C. & Golemis, E. A. Adaptors for disorders of the brain? The cancer signaling proteins NEDD9, CASS4, and PTK2B in Alzheimer’s disease. Oncoscience (2014). doi:10.18632/oncoscience.64

64. Karch, C. M. & Goate, A. M. Alzheimer’s disease risk genes and mechanisms of disease pathogenesis. Biological Psychiatry (2015). doi:10.1016/j.biopsych.2014.05.006

65. Kempson, S. A., Zhou, Y. & Danbolt, N. C. The betaine/GABA transporter and betaine: Roles in brain, kidney, and liver. Frontiers in Physiology (2014). doi:10.3389/fphys.2014.00159

66. Iadecola, C., Anrather, J. & Kamel, H. Effects of COVID-19 on the Nervous System. Cell (2020). doi:10.1016/j.cell.2020.08.028

67. Yang, A. C. et al. Broad transcriptional dysregulation of brain and choroid plexus cell types with COVID-19. bioRxiv 2020.10.22.349415 (2020). doi:10.1101/2020.10.22.349415

68. Parker, K. R. et al. Single-Cell Analyses Identify Brain Mural Cells Expressing CD19 as Potential Off-Tumor Targets for CAR-T Immunotherapies. Cell (2020). doi:10.1016/j.cell.2020.08.022

69. Månberg, A. et al. Altered perivascular fibroblast activity precedes ALS disease onset. Nat Med 27, (2021).

70. Halpern, K. B. et al. Single-cell spatial reconstruction reveals global division of labour in the mammalian liver. Nature (2017). doi:10.1038/nature21065

71. He, L. et al. Analysis of the brain mural cell transcriptome. Sci. Rep. (2016). doi:10.1038/srep35108

72. Blinder, P. et al. The cortical angiome: An interconnected vascular network with noncolumnar patterns of blood flow. Nat. Neurosci. (2013). doi:10.1038/nn.3426

73. Sabbagh, M. F. et al. Transcriptional and epigenomic landscapes of CNS and non-CNS vascular endothelial cells. Elife (2018). doi:10.7554/eLife.36187

74. Trapnell, C. et al. The dynamics and regulators of cell fate decisions are revealed by pseudotemporal ordering of single cells. Nat. Biotechnol. (2014). doi:10.1038/nbt.2859

75. Uhlén, M. et al. Tissue-based map of the human proteome. Science (80-.). (2015). doi:10.1126/science.1260419

76. De Meyer, S. F., Stoll, G., Wagner, D. D. & Kleinschnitz, C. Von Willebrand factor: An emerging target in stroke therapy. Stroke (2012). doi:10.1161/STROKEAHA.111.628867

77. Cooney, M. T., Dudina, A. L., O’Callaghan, P. & Graham, I. M. Von Willebrand factor in CHD and stroke: Relationships and therapeutic implications. Current Treatment Options in Cardiovascular Medicine (2007). doi:10.1007/s11936-007-0011-8

78. Mergenthaler, P. & Meisel, A. Do stroke models model stroke? DMM Disease Models and Mechanisms (2012). doi:10.1242/dmm.010033

79. Fluri, F., Schuhmann, M. K. & Kleinschnitz, C. Animal models of ischemic stroke and their application in clinical research. Drug Des. Devel. Ther. (2015). doi:10.2147/DDDT.S56071

80. Joutel, A., Haddad, I., Ratelade, J. & Nelson, M. T. Perturbations of the cerebrovascular matrisome: A convergent mechanism in small vessel disease of the brain? Journal of Cerebral Blood Flow and Metabolism (2016). doi:10.1038/jcbfm.2015.62

81. Mao, M., Alavi, M. V., Labelle-Dumais, C. & Gould, D. B. Type IV Collagens and Basement Membrane Diseases: Cell Biology and Pathogenic Mechanisms. Curr. Top. Membr. (2015). doi:10.1016/bs.ctm.2015.09.002

82. Zeisel, A. et al. Cell types in the mouse cortex and hippocampus revealed by single-cell RNA-seq. Science (80-.). (2015). doi:10.1126/science.aaa1934

83. Zhang, Y. et al. An RNA-sequencing transcriptome and splicing database of glia, neurons, and vascular cells of the cerebral cortex. J. Neurosci. (2014). doi:10.1523/JNEUROSCI.1860-14.2014

84. Mastorakos, P. & McGavern, D. The anatomy and immunology of vasculature in the central nervous system. Science Immunology (2019). doi:10.1126/sciimmunol.aav0492

85. Iliff, J. J. et al. A paravascular pathway facilitates CSF flow through the brain parenchyma and the clearance of interstitial solutes, including amyloid β. Sci. Transl. Med. (2012). doi:10.1126/scitranslmed.3003748

86. Louveau, A. et al. Structural and functional features of central nervous system lymphatic vessels. Nature (2015). doi:10.1038/nature14432

87. Da Mesquita, S. et al. Functional aspects of meningeal lymphatics in ageing and Alzheimer’s disease. Nature (2018). doi:10.1038/s41586-018-0368-8

88. Aspelund, A. et al. A dural lymphatic vascular system that drains brain interstitial fluid and macromolecules. J. Exp. Med. (2015). doi:10.1084/jem.20142290

89. DeSisto, J. et al. Single-Cell Transcriptomic Analyses of the Developing Meninges Reveal Meningeal Fibroblast Diversity and Function. Dev. Cell (2020). doi:10.1016/j.devcel.2020.06.009

90. Vanlandewijck, M. et al. A molecular atlas of cell types and zonation in the brain vasculature. Nature 554, 475–480 (2018).

91. Dorrier, C. E. et al. CNS fibroblasts form a fibrotic scar in response to immune cell infiltration. Nat. Neurosci. (2021). doi:10.1038/s41593-020-00770-9

92. Soderblom, C. et al. Perivascular fibroblasts form the fibrotic scar after contusive spinal cord injury. J. Neurosci. (2013). doi:10.1523/JNEUROSCI.2524-13.2013

93. Jin, S., Guerrero-juarez, C. F., Zhang, L., Chang, I. & Myung, P. Inference and analysis of cell-cell communication using CellChat. Nat. Commun. (2020).

94. Efremova, M., Vento-Tormo, M., Teichmann, S. A. & Vento-Tormo, R. CellPhoneDB: inferring cell–cell communication from combined expression of multi-subunit ligand–receptor complexes. Nat. Protoc. (2020). doi:10.1038/s41596-020-0292-x

95. Wyss-Coray, T., Lin, C., Sanan, D. A., Mucke, L. & Masliah, E. Chronic overproduction of transforming growth factor-β1 by astrocytes promotes Alzheimer’s disease-like microvascular degeneration in transgenic mice. Am. J. Pathol. (2000). doi:10.1016/S0002-9440(10)64713-X

96. Wyss-Coray, T. et al. Amyloidogenic role of cytokine TGF-β1 in transgenic mice and in Alzheimer’s disease. Nature (1997). doi:10.1038/39321

97. Masters, C. L. et al. Alzheimer’s disease. Nature Reviews Disease Primers (2015). doi:10.1038/nrdp.2015.56

98. Braak, H. & Braak, E. Neuropathological stageing of Alzheimer-related changes. Acta Neuropathologica (1991). doi:10.1007/BF00308809

99. De Strooper, B. & Karran, E. The Cellular Phase of Alzheimer’s Disease. Cell (2016). doi:10.1016/j.cell.2015.12.056

100. Heneka, M. T. et al. Neuroinflammation in Alzheimer’s disease. The Lancet Neurology (2015). doi:10.1016/S1474-4422(15)70016-5

101. Bishop, N. A., Lu, T. & Yankner, B. A. Neural mechanisms of ageing and cognitive decline. Nature (2010). doi:10.1038/nature08983

102. Keren-Shaul, H. et al. A Unique Microglia Type Associated with Restricting Development of Alzheimer’s Disease. Cell (2017). doi:10.1016/j.cell.2017.05.018

103. Zhao, Z., Nelson, A. R., Betsholtz, C. & Zlokovic, B. V. Establishment and Dysfunction of the Blood-Brain Barrier. Cell (2015). doi:10.1016/j.cell.2015.10.067

104. Sweeney, M. D., Sagare, A. P. & Zlokovic, B. V. Blood-brain barrier breakdown in Alzheimer disease and other neurodegenerative disorders. Nature Reviews Neurology (2018). doi:10.1038/nrneurol.2017.188

105. Roher, A. E. et al. Cerebral blood flow in Alzheimer’s disease. Vasc. Health Risk Manag. (2012). doi:10.2147/VHRM.S34874

106. Johnson, N. A. et al. Pattern of cerebral hypoperfusion in Alzheimer disease and mild cognitive impairment measured with arterial spin-labeling MR imaging: Initial experience. Radiology (2005). doi:10.1148/radiol.2343040197

107. Tournier-Lasserve, E. et al. Cerebral autosomal dominant arteriopathy with subcortical infarcts and leukoencephalopathy maps to chromosome 19q12. Nat. Genet. (1993). doi:10.1038/ng0393-256

108. Tikka, S. et al. CADASIL and CARASIL. Brain pathology (Zurich, Switzerland) (2014). doi:10.1111/bpa.12181

109. Kashyap, P. V., Bhat, S. J., Bhatt, S. & Dhasmana, M. Cerebral autosomal dominant arteriopathy with subcortical infarcts and leukoencephalopathy (CADASIL). J. Assoc. Physicians India (2019). doi:10.2298/vsp1105455k

110. Kapoor, A. & Nation, D. A. Role of Notch signaling in neurovascular aging and Alzheimer’s disease. Semin. Cell Dev. Biol. (2020). doi:10.1016/j.semcdb.2020.12.011

111. Montagne, A. et al. APOE4 leads to blood–brain barrier dysfunction predicting cognitive decline. Nature (2020). doi:10.1038/s41586-020-2247-3

112. Rockenstein, E., Mallory, M., Mante, M., Sisk, A. & Masliaha, E. Early formation of mature amyloid-β protein deposits in a mutant APP transgenic model depends on levels of Aβ1-42. J. Neurosci. Res. (2001). doi:10.1002/jnr.1247

113. Thal, D. R., Rüb, U., Orantes, M. & Braak, H. Phases of Aβ-deposition in the human brain and its relevance for the development of AD. Neurology (2002). doi:10.1212/WNL.58.12.1791

114. Montagne, A. et al. Blood-Brain barrier breakdown in the aging human hippocampus. Neuron 85, 296–302 (2015).

115. Maurano, M. T. et al. Systematic localization of common disease-associated variation in regulatory DNA. Science (80-.). (2012). doi:10.1126/science.1222794

116. Nott, A. et al. Brain cell type–specific enhancer–promoter interactome maps and disease-risk association. Science (80-.). (2019). doi:10.1126/science.aay0793

117. Lambert, J. C. et al. Meta-analysis of 74,046 individuals identifies 11 new susceptibility loci for Alzheimer’s disease. Nat. Genet. (2013). doi:10.1038/ng.2802

118. Kunkle, B. W. et al. Genetic meta-analysis of diagnosed Alzheimer’s disease identifies new risk loci and implicates Aβ, tau, immunity and lipid processing. Nat. Genet. (2019). doi:10.1038/s41588-019-0358-2

119. Jansen, I. E. et al. Genome-wide meta-analysis identifies new loci and functional pathways influencing Alzheimer’s disease risk. Nat. Genet. (2019). doi:10.1038/s41588-018-0311-9

120. Efthymiou, A. G. & Goate, A. M. Late onset Alzheimer’s disease genetics implicates microglial pathways in disease risk. Molecular Neurodegeneration (2017). doi:10.1186/s13024-017-0184-x

121. Hardy, J. & Escott-Price, V. Genes, pathways and risk prediction in Alzheimer’s disease. Human Molecular Genetics (2019). doi:10.1093/hmg/ddz163

122. Harold, D. et al. Genome-wide association study identifies variants at CLU and PICALM associated with Alzheimer’s disease. Nat. Genet. (2009). doi:10.1038/ng.440

123. Wightman, D. P. et al. Largest GWAS (N=1,126,563) of Alzheimer’s Disease Implicates Microglia and Immune Cells. medRxiv (2020). doi:10.1101/2020.11.20.20235275

124. Skene, N. G. & Grant, S. G. N. Identification of vulnerable cell types in major brain disorders using single cell transcriptomes and expression weighted cell type enrichment. Front. Neurosci. (2016). doi:10.3389/fnins.2016.00016

125. Hardy, J. et al. Pathways to Alzheimer’s disease. J. Intern. Med. (2014). doi:10.1111/joim.12192

126. Bero, A. W. et al. Neuronal activity regulates the regional vulnerability to amyloid-β 2 deposition. Nat. Neurosci. (2011). doi:10.1038/nn.2801

127. Safaiyan, S. et al. Age-related myelin degradation burdens the clearance function of microglia during aging. Nat. Neurosci. (2016). doi:10.1038/nn.4325

128. Spangenberg, E. et al. Sustained microglial depletion with CSF1R inhibitor impairs parenchymal plaque development in an Alzheimer’s disease model. Nat. Commun. (2019). doi:10.1038/s41467-019-11674-z

129. Huang, Y. et al. Repopulated microglia are solely derived from the proliferation of residual microglia after acute depletion. Nat. Neurosci. (2018). doi:10.1038/s41593-018-0090-8

130. Arrojo e Drigo, R. et al. Age Mosaicism across Multiple Scales in Adult Tissues. Cell Metab. (2019). doi:10.1016/j.cmet.2019.05.010

131. Nortley, R. et al. Amyloid b oligomers constrict human capillaries in Alzheimer’s disease via signaling to pericytes. Science (80-.). (2019). doi:10.1126/science.aav9518

132. Gate, D. et al. Clonally expanded CD8 T cells patrol the cerebrospinal fluid in Alzheimer’s disease. Nature (2020). doi:10.1038/s41586-019-1895-7

133. Farh, K. K. H. et al. Genetic and epigenetic fine mapping of causal autoimmune disease variants. Nature (2015). doi:10.1038/nature13835

134. Villar, D. et al. Enhancer evolution across 20 mammalian species. Cell (2015). doi:10.1016/j.cell.2015.01.006

135. Buenrostro, J. D., Wu, B., Chang, H. Y. & Greenleaf, W. J. ATAC-seq: A method for assaying chromatin accessibility genome-wide. Curr. Protoc. Mol. Biol. (2015). doi:10.1002/0471142727.mb2129s109

136. Park, P. J. ChIP-seq: Advantages and challenges of a maturing technology. Nature Reviews Genetics (2009). doi:10.1038/nrg2641

137. Fang, R. et al. Mapping of long-range chromatin interactions by proximity ligation-assisted ChIP-seq. Cell Research (2016). doi:10.1038/cr.2016.137

138. Regev, A. et al. The human cell atlas. Elife (2017). doi:10.7554/eLife.27041

139. Schirmer, L. et al. Neuronal vulnerability and multilineage diversity in multiple sclerosis. Nature (2019). doi:10.1038/s41586-019-1404-z

140. Thul, P. J. et al. A subcellular map of the human proteome. Science (80-.). (2017). doi:10.1126/science.aal3321

141. Zhou, Y. et al. Metascape provides a biologist-oriented resource for the analysis of systems-level datasets. Nat. Commun. (2019). doi:10.1038/s41467-019-09234-6

142. McInnes, L., Healy, J., Saul, N. & Großberger, L. UMAP: Uniform Manifold Approximation and Projection. J. Open Source Softw. (2018). doi:10.21105/joss.00861

143. Satija, R., Farrell, J. A., Gennert, D., Schier, A. F. & Regev, A. Spatial reconstruction of single-cell gene expression data. Nat. Biotechnol. (2015). doi:10.1038/nbt.3192

144. McGinnis, C. S., Murrow, L. M. & Gartner, Z. J. DoubletFinder: Doublet Detection in Single-Cell RNA Sequencing Data Using Artificial Nearest Neighbors. Cell Syst. (2019). doi:10.1016/j.cels.2019.03.003

145. Finak, G. et al. MAST: A flexible statistical framework for assessing transcriptional changes and characterizing heterogeneity in single-cell RNA sequencing data. Genome Biol. (2015). doi:10.1186/s13059-015-0844-5

146. Lake, B. B. et al. Integrative single-cell analysis of transcriptional and epigenetic states in the human adult brain. Nat. Biotechnol. (2018). doi:10.1038/nbt.4038

147. Chen, E. Y. et al. Enrichr: Interactive and collaborative HTML5 gene list enrichment analysis tool. BMC Bioinformatics (2013). doi:10.1186/1471-2105-14-128

148. Conway, J. R., Lex, A. & Gehlenborg, N. UpSetR: An R package for the visualization of intersecting sets and their properties. Bioinformatics (2017). doi:10.1093/bioinformatics/btx364

149. Swiech, L. et al. In vivo interrogation of gene function in the mammalian brain using CRISPR-Cas9. Nat. Biotechnol. (2015). doi:10.1038/nbt.3055

150. Butler, A., Hoffman, P., Smibert, P., Papalexi, E. & Satija, R. Integrating single-cell transcriptomic data across different conditions, technologies, and species. Nat. Biotechnol. (2018). doi:10.1038/nbt.4096

151. Buniello, A. et al. The NHGRI-EBI GWAS Catalog of published genome-wide association studies, targeted arrays and summary statistics 2019. Nucleic Acids Res. (2019). doi:10.1093/nar/gky1120

152. Chen, M. B. et al. Brain Endothelial Cells Are Exquisite Sensors of Age-Related Circulatory Cues. Cell Rep. (2020). doi:10.1016/j.celrep.2020.03.012

153. Yang, A. C. et al. Multiple Click-Selective tRNA Synthetases Expand Mammalian Cell-Specific Proteomics. J. Am. Chem. Soc. (2018). doi:10.1021/jacs.8b03074

154. Szabo, P. A. et al. Single-cell transcriptomics of human T cells reveals tissue and activation signatures in health and disease. Nat. Commun. (2019). doi:10.1038/s41467-019-12464-3

155. Baeten, K. M. & Akassoglou, K. Extracellular matrix and matrix receptors in blood-brain barrier formation and stroke. Dev. Neurobiol. (2011). doi:10.1002/dneu.20954

